# Loss of flight in the Galapagos cormorant mirrors human skeletal ciliopathies

**DOI:** 10.1101/061432

**Authors:** Alejandro Burga, Weiguang Wang, Paul C. Wolf, Andrew M. Ramey, Claudio Verdugo, Karen Lyons, Patricia G. Parker, Leonid Kruglyak

## Abstract

Changes in the size and proportion of limbs and other structures have played a key role in the adaptive evolution of species. However, despite the ubiquity of these modifications, we have a very limited idea of how these changes occur on the genetic and molecular levels. To fill this gap, we studied a recent and extreme case of wing and pectoral skeleton size reduction leading to flightlessness in the Galapagos cormorant (*Phalacrocorax harrisi*). We sequenced and de novo assembled the genomes of four closely related cormorant species and applied a joint predictive and comparative genomics approach to find candidate variants. Here we report that function-altering variants in genes necessary for both the correct transcriptional regulation and function of the primary cilium contributed to the evolution of loss of flight in *P. harrisi*. Cilia are essential for Hedgehog signaling, and humans affected by skeletal ciliopathies suffer from premature arrest of bone growth, mirroring the skeletal features associated with loss of flight.

## Introduction

The evolution of loss of flight is one the most recurrent limb modifications encountered in nature. In fact, Darwin used the occurrence of flightless birds as an argument in favor of his theory of natural selection (Darwin, 1859). He proposed that loss of flight could evolve by two mechanisms: 1) selection in favor of larger bodies and 2) relaxed selection due to the absence of predators. Loss of flight has evolved repeatedly, and cases are found among 26 families of birds in 17 different orders (Roff, 1994). Moreover, recent studies strongly suggest that the ratites (ostriches, emus, rheas, cassowaries and kiwis), long thought to derive from a single flightless ancestor, actually constitute a polyphyletic group characterized by multiple independent instances of loss of flight and convergent evolution (Baker et al., 2014; Harshman et al., 2008; Mitchell et al., 2014). However, despite the ubiquity and evolutionary importance of loss of flight (Wright et al., 2016), the underlying genetic and molecular mechanisms remain largely unknown.

The Galapagos cormorant (*Phalacrocorax harrisi*) is the only flightless cormorant among approximately 40 extant species (Livezey, 1992). The entire population is distributed along the coastlines of Isabela and Fernandina Islands in the Galapagos archipelago. *P. harrisi* has a pair of short but perfectly shaped wings, which are smaller than those of any other cormorant (Fig. 1a). This results in a significant deviation from the well-established allometric relationship between wing length and body mass (Livezey, 1992). In addition, the Galapagos cormorant has a highly reduced keel and underdeveloped pectoral muscles. The keel or carina is an extension of the sternum that runs along its midline and provides an attachment surface for the flight muscles, the largest muscles in birds. Flightless taxa, such as ratites and Cretaceous *Hesperornis*, typically evolve flat sternums in which the keel has been largely reduced or lost (Feduccia, 1999). Surprisingly, despite its striking phenotype, the Galapagos Cormorant escaped Darwins attention during his discovery voyage on the Beagle. In contrast to ratites and penguins, which became flightless over 50 million years ago (MYA) (Mitchell et al., 2014; Slack et al., 2006), the Galapagos cormorant and its flighted relatives are estimated to share a common ancestor 2 MYA (Kennedy et al., 2009). This recent and extreme modification of wing size and pectoral skeleton makes *P. harrisi* an attractive model to study loss of flight.

To investigate the genetics of flightlessness evolution, we applied a joint predictive and comparative genomics approach to first identify all coding variants between the Galapagos cormorant and its flighted relatives, and to then classify these variants according to their probability of affecting protein function. Among the strongest function-altering variants present in *P. harrisi*, we found a significant enrichment for genes mutated in human skeletal ciliopathies. The primary cilium is essential for Hedgehog (Hh) signaling in vertebrates, and individuals affected by ciliopathies have small limbs and rib cages, mirroring the phenotype of *P. harrisi*. Our results indicate that the combined effect of variants in genes necessary for both the correct transcriptional regulation and function of the primary cilium contributed to the evolution of highly reduced wings and other skeletal adaptations associated with loss of flight in *P. harrisi*.

### High quality genome sequences of four cormorant species

To identify variants associated with loss of flight, we sequenced and de novo assembled the 1.2 Gb genomes of the Galapagos cormorant (Galapagos Islands, Ecuador) and three flighted cormorant species: the Doublecrested cormorant (*Phalacrocorax auritus*; Minnesota, USA), the Neotropical cormorant (*Phalacrocorax brasilianus*; Valdivia, Chile), and the Pelagic cormorant (*Phalacrocorax pelagicus*; Alaska, USA) (Supplementary Fig. 1). *P. auritus* and *P. brasilianus* are the closest relatives of *P. harrisi* (Kennedy et al., 2009), and P. pelagicus is part of a sister clade and served as an outgroup. Genomes were assembled from a combination of short insert and mate-pair Illumina libraries using SOAPdenovo2 (Luo et al., 2012). Details about the sequenced individuals and library statistics can be found in Supplementary Table 1a and Supplementary Table 2. Among these four genomes, the Galapagos cormorants assembly had the longest contig and scaffold N50 metrics (contig N50: 103 kb; scaffold N50: 4.6 Mb; Supplementary Table 1b). We evaluated the completeness of the cormorants genomes by estimating the total number of uniquely annotated proteins in each assembly and by using the CEGMA pipeline (Parra et al., 2009)(see Methods). Overall, we found good agreement between these two independent metrics in a dataset including the four cormorant genomes and 17 recently published bird genomes (r^2^ = 0.75 P = 4.3×10-07; Fig. 1b and Supplementary Table 3). Interestingly, commonly used metrics of assembly quality such as contig and scaffold N50 were very poor predictors of the total number of proteins present in each assembly (r^2^ = 0.13 P = 0.15 and r^2^ = 0.06 P = 0.81; Supplementary Fig. 2 Supplementary Table 3).Three of the four cormorant genomes (*P. harrisi*, textitP. auritus and textitP. pelagicus) obtained the highest CEGMA scores and number of uniquely annotated genes of all bird genomes (red triangles, Fig. 1d). For instance, the following CEGMA scores were obtained for the cormorants: *P. harrisi*, 90.3%; *P. auritus*, 91.3%; *P. brasilianus*, 72.6%; *P. pelagicus*, 87.1%. In contrast, Sanger and PacBio genomes obtained lower scores: *Gallus gallus* (Sanger assembly) (Consortium, 2004), 80.7%; *Taeniopygia guttata* (Sanger assembly)(Warren et al., 2010), 71.4%; and *Melopsittacus undulatus* (PacBio assembly) (Ganapathy et al., 2014), 79.0%. Thus, our cormorant genomes are the most complete bird genomes published to date from a gene-centric perspective, performing even better than genomes assembled from Sanger sequences and PacBio long reads (complete statistics in Supplementary Table 3).

**Figure 1:**
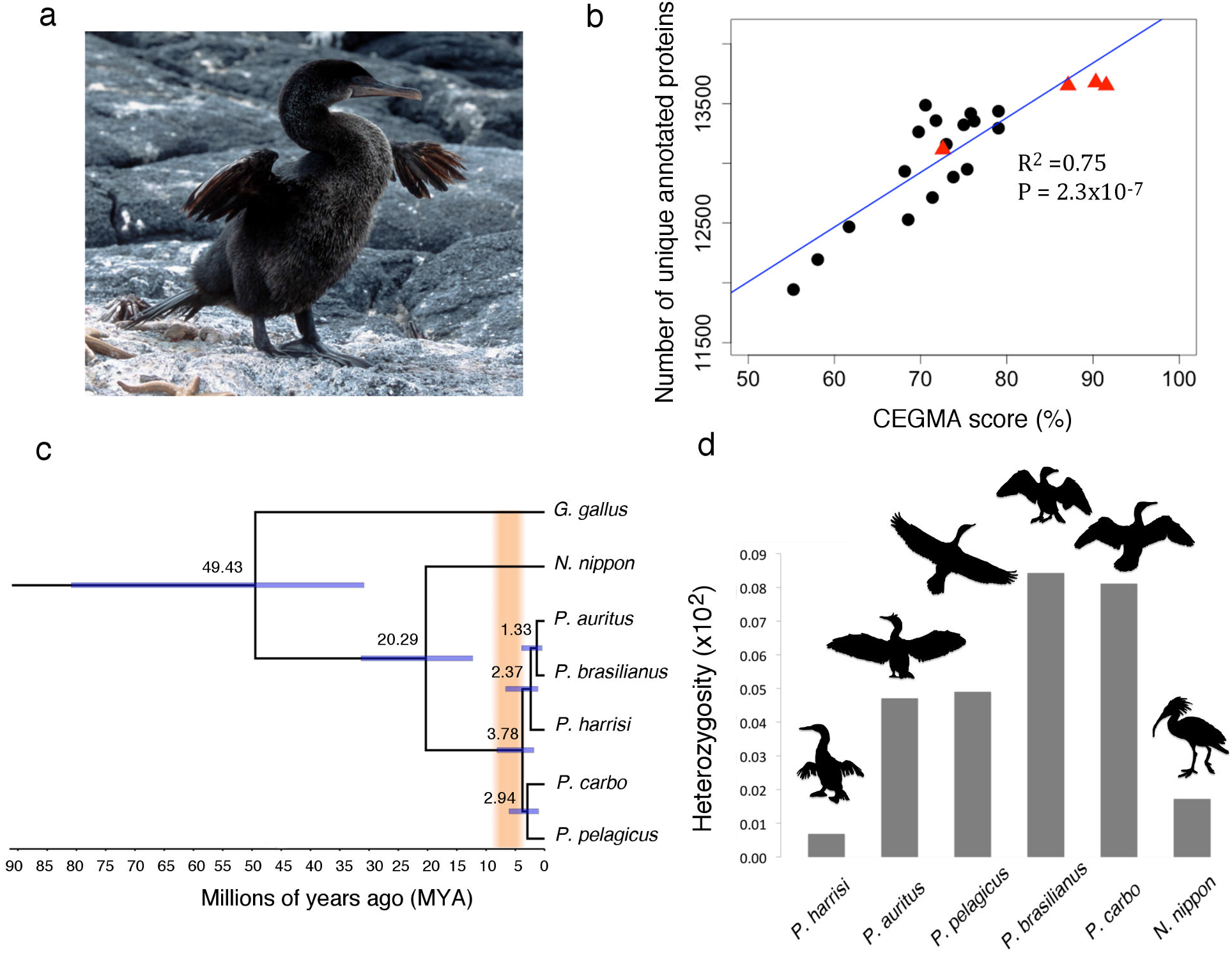
The Galapagos Cormorant, a model to study flightlessness evolution. a, The Galapagos cormorant has evolved very short wings. The average wing length of an adult male is 19 cm (3.6 kg. body mass), whereas the wing length of its closest relative, the doublecrested cormorant, is 31.5 cm. (2.2 kg. body mass). b, CEGMA score is a good predictor of genome completeness from a gene-centric perspective. Blue line is the linear regression model (r^2^ = 0.75 P = 4.3×10-07). Genomes reported in this study are red triangles and other published avian genomes are black circles (see Supplementary Table 3 for full details). c, Phylogenetic tree reconstructed using fourfold degenerate sites from whole genome sequences. In orange, time span between approximate origin of proto-Galapagos archipelago (9 MYA) and the origin of the oldest extant island, San Cristobal (4 MYA). Nodes represent median divergence ages. Blue bars indicate the 95% Highest Posterior Density (HPD) Interval. d, Heterozygosity levels inferred from whole genome sequences. Birds are not drawn to scale.

### Phylogeny and genetic diversity

We reconstructed the cormorant phylogeny using a Bayesian framework (see Methods). Our analysis confirmed the previously reported phylogenetic relationships among the four sequenced species (Fig. 1c). Moreover, our results indicate that *P. harrisi* last shared a common ancestor with *P. auritus* and *P. brasilianus* 2.37 MYA, in agreement with a previous estimate based on mitochondrial DNA (Kennedy et al., 2009) (Fig. 1c). The oldest extant island in the Galapagos archipelago, Espaola, emerged at most 4 MYA, and proto-Galapagos islands existed at least 9 MYA (Harpp, 2014). Our results are consistent with the view that *P. harrisi* lost the ability to fly while inhabiting the archipelago. We calculated the proportion of single nucleotide polymorphism (SNP) heterozygous sites for each sequenced individual to estimate the levels of intra-specific genetic diversity (Fig. 1d). *P. harrisi* showed the lowest proportion of heterozygous SNPs among the sequenced cormorants (0.00685%; Fig. 1d). The heterozygosity of *P. harrisi* is even lower than that of the Crested Ibis, *Nipponia nippon*, a highly endangered bird that nearly went extinct in the 1980s (Wang and Li, 2008) (0.0172%; Fig. 1d). The low level of heterozygosity found in the Galapagos cormorant is most likely due to its small population size ( 1,500 individuals) and multiple population bottlenecks (Valle and Coulter, 1987).

### Discovery and characterization of function-altering variants in *P. harrisi*

For our variant discovery approach to be effective, it was imperative to interrogate most of the Galapagos cormorants genes. To increase our power to do so, we annotated genes using two different strategies: homology-based and transcriptome-based gene annotations (Supplementary Fig. 3). The latter was derived using mRNA expression data from the developing wing of a double-crested cormorant embryo (Supplementary Fig. 5c, see Methods). We then predicted all missense, deletion, and insertion variants in ortholog pairs between *P. harrisi* and each of its three flighted relatives (see Methods and Supplementary Fig. 2). We used PROVEAN (Choi et al., 2012), a phylogeny-corrected variant effect predictor, to evaluate the impact on protein function of each of the Galapagos cormorants variants on a genome-wide scale. A PROVEAN score is calculated for each variant; the more negative the score, the more likely a given variant is to affect protein function in the Galapagos cormorant. To illustrate our approach, Fig. 2a shows the distribution of PROVEAN scores obtained when comparing 12,442 ortholog pairs between *P. harrisi* and *P. auritus*. Of these 12,442 ortholog pairs, 4,959 (40%) did not contain coding variants; the remaining 7,483 pairs contained a total of 23,402 coding variants. These variants were further classified as: 22,643 single amino acid substitutions, 456 deletions, and 303 insertions (Fig. 2b). Most variants were predicted to be neutral (distribution is centered around zero). As expected, deletion and insertions were enriched in the tails of the distributions (Fig. 2b). Very similar numbers of variants and PROVEAN score distributions were obtained for the other homology-based (Supplementary Fig. 4) and transcriptome-based annotations (Supplementary Fig. 5).

### Enrichment for genes mutated in skeletal ciliopathies

We reasoned that variants responsible for loss of flight should be enriched among those affecting protein function in *P. harrisi*. To find those variants, we applied a stringent threshold to our four prediction datasets: PROVEAN score < −5, two times the threshold for human disease variants discovery (Choi et al., 2012) (Fig. 2a, see Methods). Variants with a PROVEAN score < −5 are typically at residues that have been perfectly conserved at least since mammals and birds last shared a common ancestor (300 MYA), and consequently, they very are likely to be important for protein function.

Gene enrichment analysis of function-altering variants in the Galapagos cormorant revealed a significant overrepresentation of human developmental disorders (see Methods and Supplementary Table 4a). Strikingly, 8 out of the 19 significant categories were related to syndromes affecting limb development, such as polydactyly, syndactyly, and duplication of limb bones. Control enrichment analysis didnt reveal any limb syndrome disorder in the other flighted cormorants (see Methods, Supplementary Table 4b and 4c).

The genes underlying the enrichment for limb syndromes were, upon examination, all mutated in a family of human disorders known as ciliopathies. Cilia are hair-like microtubule based structures that are nucleated by the basal body (centriole and associated proteins) and project from the surface of cells. They are essential for mediating Hedgehog (Hh) signaling, serving as antennae for morphogens during development (Goetz and Anderson, 2010). We confirmed by Sanger sequencing the presence of function-altering variants in *Ofd1, Evc, TalpidS, Dync2h1, Ift122*, Wdr34, and Kif7, all of which are necessary for the assembly or functioning of the primary cilium and are mutated in human ciliopathies, particularly those affecting the skeleton (Table 1). Individuals affected by skeletal ciliopathies have small limbs and rib cages, mirroring the main features of the Galapagos cormorant. Furthermore, our enrichment analysis also revealed the presence of a function-altering variant in *Gli2*, a transcription factor essential for Hh signaling (Haycraft et al., 2005) (Table 1).

**Figure 2:**
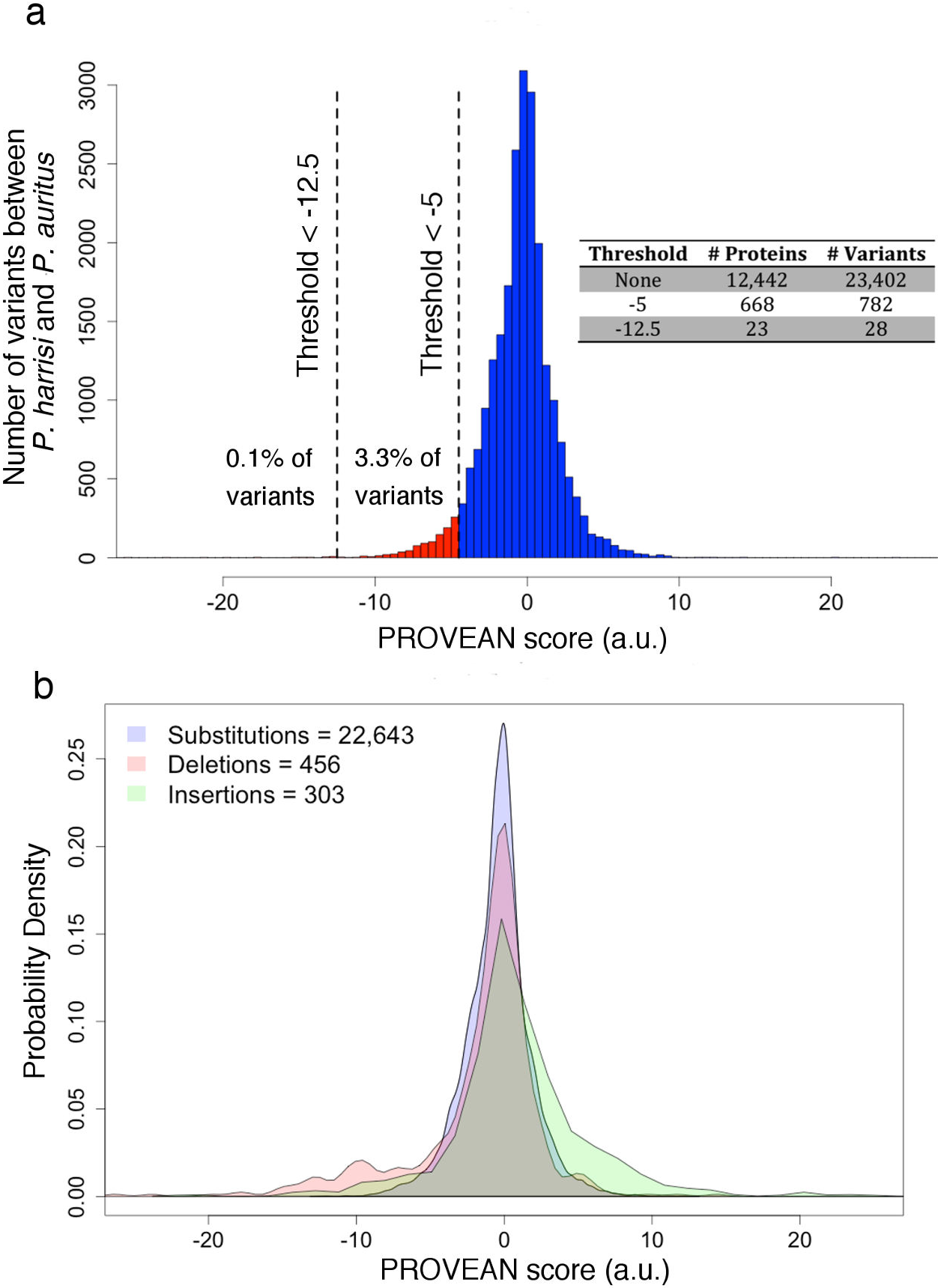
Distribution of the effect of variants between*P. auritus*. and*P. harrisi*. a, We used PROVEAN to predict the effect on protein function of 23,402 variants contained in 12,449 orthologous pairs between*P. auritus*. and*P. harrisi*. 4,966 pairs contained no variants. The more negative the score; the more likely the variant affects protein function. PROVEAN score thresholds used in this study are drawn as vertical dashed lines. Number of proteins and variants found for each threshold are presented in inset table. b, Density of PROVEAN scores for each class of variant. The same variants presented in (a) were classified as single amino acid substitutions, deletions, and insertions. Number of variants in each class is indicated in the legend.

**Table 1:** Function-altering variants in*P. harrisi* are enriched for genes that cause skeletal ciliopathies in humans. Sanger validated examples of function-altering variants (PROVEAN score < −5) in P.harrisi. Cilia/Hh related genes were found based on functional enrichment for human syndromes. PCP (Planar cell polarity) genes were selected based on literature evidence linking cilia and PCP. These variant are fixed in the population. (*) Based on phenotypic similarity to mutant mouse model

| Gene | Pathway | Variant | PROVEAN score | Human Syndrome |
| --- | --- | --- | --- | --- |
| *Ofd1* | Cilia/Hh | R325C | -6.913 | Orofaciodigital and Joubert |
|  |  | K517T | -5.673 |  |
|  |  | E889G | -5.068 |  |
| *Talpid3* | Cilia/Hh | D759V | -7.805 | Joubert and Jeune |
| *Evc* | Cilia/Hh | T341I | -5.546 | Ellias-Van Creveld |
| *Dync2h1* | Cilia/Hh | P2733S | -7.431 | Short-rib thoracic dysplasia |
| *Ift122* | Cilia/Hh | Q691L | -5.491 | Cranioectodermal dysplasia |
| *Wdr34* | Cilia/Hh | P188R | -6.337 | Short-rib thoracic dysplasia |
| *Kif7* | Cilia/Hh | R833W | -6.827 | Joubert and Acrocallosal |
| *Gli2* | Hh | P1086T | -5.117 | Culler-Jones |
| *Fat1* | PCP | S1717L | -5.858 | Faciocapulohumeral Dystrophy (*) |
|  |  | Y2462C | -8.592 |  |
| *Dchs1* | PCP | G2063D | -6.45 | Van Maldergem |
| *Dvl1* | PCP | P103L | -8.23 | Robinow |

The Galapagos cormorants ortholog of human OFD1 (mutated in Oral-facial-digital syndrome 1) is affected by three function-altering variants with PROVEAN score < −5 (R325C −6.913, K517T −5.673 and E889G −5.068; Supplementary Fig. 6a and 6b). *Ofd1* mutant mice display polydactyly and shortened long bones (Bimonte et al., 2011). Also, a function-altering missense variant (Q691L −5.491) was found in *Ift122*, a component of the IFT complex that controls the ciliary localization of Shh pathway regulators (Ocbina et al., 2011). The mutated glutamine in *Ift122* is virtually invariant among eukaryotes ranging from green algae to vertebrates (Supplementary Fig. 6c).

There is a well-established genetic link between the Planar Cell Polarity (PCP) pathway and cilia (Goetz and Anderson, 2010). For instance, intu and fuz are core downstream effectors of the PCP pathway in Drosophila, and their respective Xenopus (Park et al., 2006) and mouse (Gray et al., 2009; Zeng et al., 2010) mutants are characterized by short cilia and disrupted Hh signaling. We searched our *P. harrisi* data set (PROVEAN score < −5) for members of the PCP pathway and found function-altering variants in *Fat1* atypical cadherin (Fat1), Dachsous cadherin-related 1 (Dchs1) and Disheveled-1 (*Dvl1*) (Supplementary Fig. 6d; Table 1). The Galapagos Cormorants FAT1 contains two function-altering variants (S1717L and Y2462C, Table 1) and *Fat1* mouse mutants show very selective defects in muscles of the upper body (Caruso et al., 2013). In addition, *Dvl1* is mutated in patients with Robinow syndrome, characterized by limb shortening (Bunn et al., 2015; White et al., 2015).

Additional Sanger sequencing of 20 Galapagos cormorant individuals from two different populations (Cabo Hammond and Caones Sur (Duffie et al., 2009)) revealed only homozygous carriers for all of the variants in Table 1, indicating that these variants are most likely fixed in the Galapagos Cormorant. Overall, we found an overrepresentation of function-altering variants in genes that, when mutated in humans, cause skeletal ciliopathies and bone growth defects.

### Cuxl is mutated in *P. harrisi.*

To identify the strongest function-altering variants in*P. harrisi*, we applied a more stringent PROVEAN score threshold: −12.5 delta alignment score, five times the threshold used for human disease variants discovery (Choi et al., 2012). This strategy narrowed our search to 23 proteins. These represent 0.16% of annotated proteins in*P. harrisi* (Supplementary Table 5). We manually curated these 23 proteins and performed additional Sanger sequencing reducing the list of proteins with confirmed or putative variants to 12. (Supplementary Table 5; see Supplementary Note for a detailed analysis). These variants were exclusively small deletions. Among these 12 proteins, two stood out based on their known role in development: LGALS-3 and CUX1. LGALS-3 is affected by a 7 amino acid deletion in*P. harrisi* (PROVEAN score −26.319). LGALS-3 (*Galectin-3*) is localized at the base of the primary cilium and is necessary for correct ciliogenesis in mice (Koch et al., 2010), yet it has not been implicated in human ciliopathies. Moreover, LGALS-3 physically interacts with SUFU, an important regulator of mammalian Hh signaling (Paces-Fessy et al., 2004), and knockout mice show pleiotropic defects in chondrocyte differentiation (Colnot et al., 2001).

In addition, we found a 4 amino acid deletion (PROVEAN score −15.704) in CUX1. CUX1 (cut-like homeobox 1), also known as CDP, is a highly conserved transcription factor with diverse developmental roles that contains four DNA binding domains: three CUT domains (CR1-3) and one homeodomain (HD) (Fig. 3a) (Sansregret and Nepveu, 2008). It is noteworthy that cut, the Drosophila ortholog of *Cux1*, is necessary for proper development of wings and flight muscles in flies(Sudarsan et al., 2001). Moreover, the expression pattern of cut is lost in the wing imaginal disc of flightless ant populations compared to their flighted relatives (Fave et al., 2015). In chicken, *Cux1* mRNA expression in the limb is restricted to the ectoderm bordering the Apical Ectodermal Ridge (AER) at embryonic stage 23 (Tavares et al., 2000). The AER is one of the key signaling centres that drive limb development. Furthermore, adenovirus-mediated overexpression of a dominant negative form of CUX1 in the developing chicken wing results in AER disruption and limb truncations most strongly affecting distal skeletal elements (digits, radius and ulna) (Tavares et al., 2000). Interestingly, in*P. harrisi*, the radius and ulna are disproportionately small compared to the humerus (Livezey, 1992).

We Sanger-sequenced and confirmed the predicted *Cux1* 12 base pair (bp) deletion in *P. harrisi*. We also confirmed that this variant was fixed in the population and absent in the other cormorant species (Supplementary Fig. 7a). The 12 bp deletion in *Cux1* removes four amino acids, AGSQ, immediately adjacent to the C-terminal end of the homeodomain (Fig. 3b). We will refer to this variant as CUX1-Δ4aa. Alignment of CUX1 orthologs from available vertebrate genomes revealed that the 4 missing residues are extremely conserved among tetrapods (Fig. 3b). The deleted serine is phosphorylated in human cells (Matsuoka et al., 2007), but the molecular consequences of this modification are unknown. The *Cux1* deletion does not include any of the predicted residues responsible for DNA contact and recognition (Iyaguchi et al., 2007), but given its close proximity to the homeodomain, we decided to test whether the DNA binding activity of CUX1 was affected. We performed Electrophoretic Mobility Shift Assay (EMSA) with purified CR3HD CUX1-Ancestral and CUX1-Δ4aa protein variants (Supplementary Fig. 7b), as previously described (Moon et al., 2000), and found that the DNA binding activity was not abolished in the deletion variant (Supplementary Fig. 7c). CUX1 is able to both directly repress and activate gene expression through its C-terminal tail (Mailly et al., 1996; Truscott et al., 2007). We performed a luciferase reporter assay (Mailly et al., 1996; Nishio and Walsh, 2004) and found that both variants were equally capable of repressing the expression of a UAS/tk luciferase reporter (Fig. 3d). Thus, the Galapagos cormorant CUX1-Δ4aa variant affects neither the DNA binding activity in vitro nor the C-terminal mediated repression activity in COS-7 cells.

### CUX1 regulates the expression of cilia and PCP genes

We hypothesized that the *Cux1* deletion variant is mechanistically related to the enrichment of function-altering variants in ciliopathy-related genes based on the following observations: First, transgenic mice overexpressing the CUX1-CR3HD isoform develop polycystic kidneys. Cilia from the cystic epithelial cells from these animals were twice as long as the ones derived from control epithelial cells (Cadieux et al., 2008). Second, the CUX1-CR2CR3HD isoform has been shown to directly up-regulate the expression of Ftm, a component of the cilia basal body that is mutated in human ciliopathies and involved in Shh signaling (Stratigopoulos et al., 2011). Third, *Cux1* knockout mice show deregulation of SHH expression in hair follicles (Ellis et al., 2001).

To test whether *Cux1* could globally regulate the expression of cilia genes, we analyzed expression array data from a study that profiled Hs578t cells stably expressing an shRNA against *Cux1*, as well as cells overexpressing the human CUX1-CR2CR3HD isoform (Vadnais et al., 2013). In concordance with the well-established role of *Cux1* as a regulator of cell growth and proliferation (Ramdzan and Nepveu, 2014), genes significantly up-or down-regulated in both conditions (p < 0.05 and > 1.1 fold change) were enriched for pathways such as Cell cycle and Mitotic G1-G1/S phases (P = 3.99×10-5 and 0.016, respectively: Supplementary Table 6). Importantly, we also found and enrichment for cilia-related categories such as Assembly of the primary cilium and Intraflagellar transport (P = 1.2×10-3 and 5.7×10-3 respectively; Supplementary Table 6). These results suggest that cilia-related genes are indeed enriched among *Cux1* target genes.

To further test whether *Cux1* can regulate ciliary genes in an appropriate cellular context, we generated ATDC5 stable lines expressing N-terminal HIS-tagged versions of CR3HD CUX1-Ancestral and CUX1-Δ4aa variants. ATDC5 is a well-characterized mouse chondrogenic cell line that largely recapitulates in vitro the differentiation landmarks of chondrocytes (Shukunami et al., 1996). We chose to express the CR3HD isoform because western blot analysis revealed that this was the most abundant CUX1 isoform expressed in the developing wing of mallard embryos ( 50 kDa; Fig. 3a). We performed quantitative reverse transcription PCR (RT-qPCR) on a selected number of genes containing predicted strong function-altering variants in*P. harrisi* (Table 1) and showing detectable levels of expression in ATDC5 cells. In addition, we measured the expression of *Ptch1*, the receptor of the Hh pathway. Our experiments revealed that the CUX1-Ancestral variant transcriptionally up-regulated the expression of *Ofd1* (1.7 fold, P = 1.2×10-6; Fig. 3c) and *Fat1* (1.8 fold, P = 0.029; Fig 3c) and downregulated the expression of *Ift122* (0.77 fold, P=2. 5x10-3; Fig 3c) and *Ptch1* (0.53 fold, P=1.4×10-2;Fig 3c) compared to the control line. In contrast, neither *Dync2h1* nor *Wdr34* expression levels were changed by CUX1-Ancestral overexpression (Fig. 3c). These results confirmed that cilia and Hh related genes are indeed transcriptional targets of CUX1 in chondrocytes.

**Figure 3:**
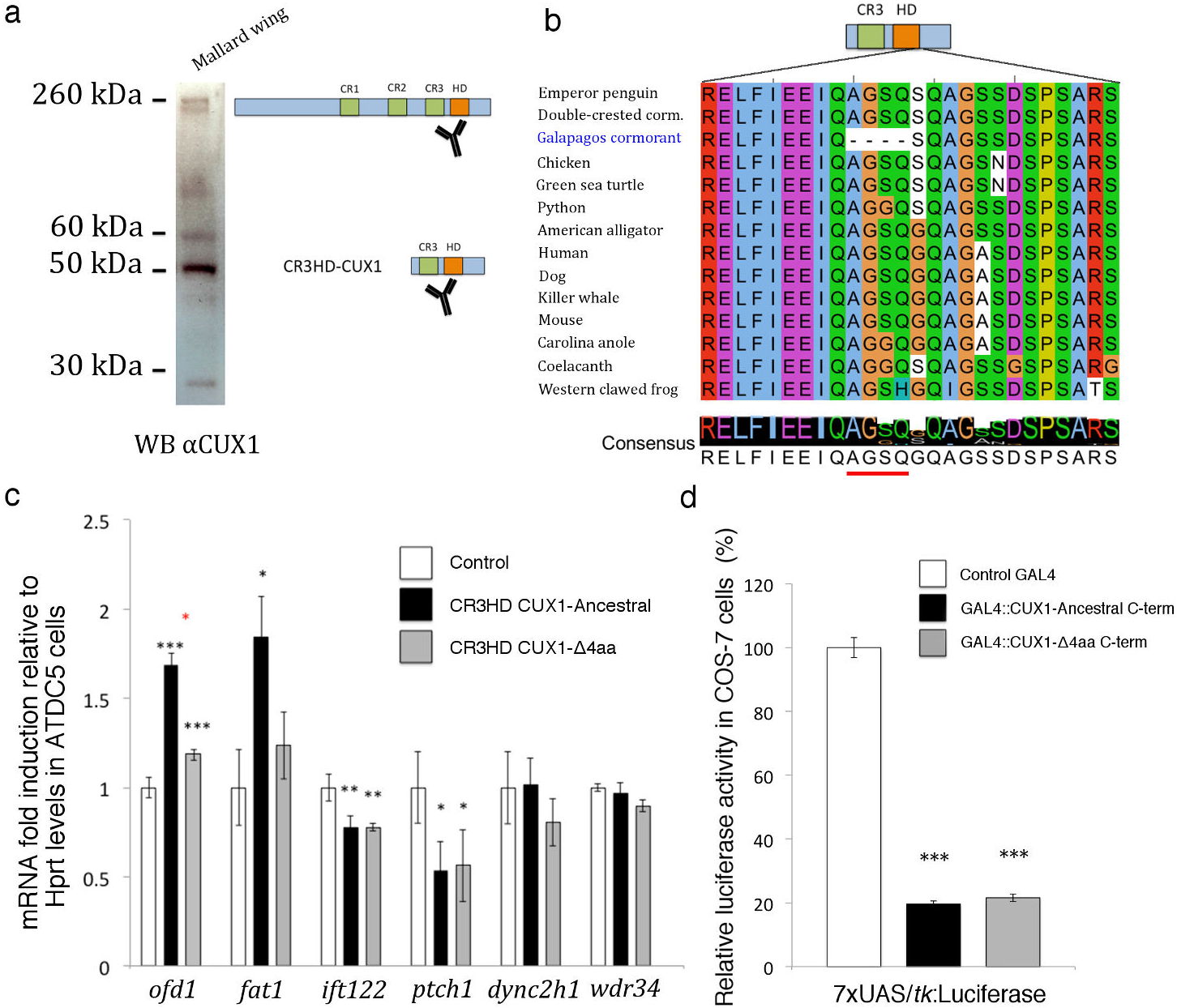
The Galapagos Cormorants Cux1 is a transcriptional activation hypomorph. a, Western blot showing the expression of CUX1 isoforms in the developing wing of a mallard embryo (22 days). The most abundant band corresponds to the predicted size of the CR3HD CUX1 isoform. b, Protein alignment showing the deleted AGSQ-residues in the Galapagos Cormorant CUX1 and their high degree of conservation among vertebrates. c, Differential up-regulation of genes by CUX1-Ancestral and CUX1-Δ4aa variants in ATDC5 cells. Pooled stable lines carrying CR3HD CUX1-Ancestral or CUX1-Δ4aa variants were generated by lentiviral transduction and puromycin selection. Controls cells were transduced with an empty vector. Gene expression levels were measured by reverse transcriptase quantitative PCR (n=5 biological replicates, each compromising 3 technical replicates). d, Luciferase based assay to test the repression activity of CUX1 C-terminal domain lacking CR3 and HD domains. GAL4 DNA binding domain was fused to CUX1-Ancestral or CUX1-Δ4aa variants. Both constructs equally repressed a promoter containing UAS binding sites in COS-7 cells (n=3 biological replicates, each compromising 3 technical replicates). Gene expression levels were measured by RT-qPCR (n=5 biological replicates, each compromising 3 technical replicates). Error bars indicate standard errors. We used ANOVA and Tukeys HSD to test for statistical significance. Black stars indicate a significant difference between a CUX1 variant and the control. Red starts indicate a significant difference between CUX1-Ancestral and CUX1-Δ4aa variants. Absence of stars indicates no significant difference. (*) P<0.05, (**) P<0.01, and (***) P<0.001

### Impaired transcriptional activity of the Galapagos cormorant CUX1

Remarkably, we found that the Galapagos cormorants CUX1 showed impaired transcriptional activity compared to the ancestral variant. *Ofd1* upregulation was significantly reduced in CUX1-Δ4aa cells compared to CUX1-Ancestral cells (1.7 vs. 1.2 fold, P = 6×10-5; Fig. 3c). Similarly, *Fat1* expression levels in CUX1-Δ4aa cells were not significantly different from control lines (1.2 fold, P=0.17; Fig 3c). However, downregulated genes (*Ift122* and *Ptch1*) showed no significant differences between CUX1-Ancestral and CUX1-Δ4aa cells (0.78 fold vs. 0.78 fold; P = 0.99 and 0.53 fold vs. 0.56 fold, P=0.96; Fig. 3c). These results indicate that the four amino acids deletion in the Galapagos Cormorant CUX1 exclusively affects its ability to activate gene expression and are consistent with our luciferase reporter assays, which show no effect on repression (Fig. 3d). Furthermore, these results indicate that both the transcriptional activator (*Cux1*) and its target genes (*Ofdl* and *Fatl*) are affected by strong function-altering variants in the Galapagos Cormorant.

### CR3HD-CUX1 promotes chondrogenesis

Chondrocytes are the main engine of bone growth. The growth of skeletal elements depends on the precise regulation of chondrocyte proliferation and hypertrophy. Mutations that affect cilia result in the premature arrest of bone growth due defects in Indian Hedgehog (IHH) signaling in chondrocytes (Kronenberg, 2003). To test the role of CUX1-CR3HD in chondrogenesis, we differentiated control, CUX1-Ancestral and CUX1-Δ4aa ATDC5 cell lines and quantified the expression of *Ihh* and *Sox9*, two well-established markers of chondrocyte differentiation in vitro and in vivo (Kronenberg, 2003). We found that overexpression of both CR3HD CUX1-Ancestral and CUX1-Δ4aa variants significantly promoted chondrogenic differentiation of ATDC5 cells after 7 and 12 days of differentiation (Fig. 4a). However, the CUX1-Δ4aa variant was not as efficient as the Ancestral variant, showing significant differences from CUX1-Ancestral in *Ihh* expression after 7 days of differentiation ( 50% decrease, P = 5.9×10-4; Fig. 4a) and for *Sox9* after 12 days ( 15% decrease, P = 1.6×10-2; Fig. 4a).

These results indicate that the Galapagos cormorant CUX1 is not as effective as the ortholog from its flighted relatives in promoting chondrogenic differentiation, and that mutations in Cux1 are sufficient to affect the dynamics of chondrogenesis. This observation is further supported by the fact that CUX1 is expressed in the hypertrophic chondrocytes of developing bones in mice, and that the bones of Cux1 mutant mice are thin and flaky (Sinclair et al., 2001)

### Possible evolutionary scenarios

Loss of flight has traditionally been attributed to relaxed selection. In such a scenario, the first cormorants that inhabited the Galapagos Islands found a unique environment that lacked predators and provided food year long, drastically reducing the need to migrate. However, we found no evidence for pseudogenization of developmental genes in*P. harrisi* (see Supplementary Note for a detailed analysis on missing genes and premature stop codons). On the other hand, loss of flight in the Galapagos Cormorant is thought to confer an advantage for diving by decreasing buoyancy (shorter wings) and indirectly allowing an increase in body size (larger oxygen storage) (Watanabe et al., 2011). This advantage could make flightlessness a target of positive selection. To evaluate if any of our candidate genes (Table 1) showed signatures of positive selection in the Galapagos Cormorant lineage, we estimated their ratio of non-synonymous to synonymous substitutions (*ω* = dN/dS). This is a very stringent selection test because it assumes all sites in a protein are evolving under the same selective pressure, a condition that is rarely met in highly conserved regulatory genes (Yang, 1997). We found that three out of eleven tested genes showed signs of positive selection (*ω* >1) in the Galapagos cormorant lineage compared to a background phylogeny of 35 Taxa (*Ofd1 ω*=1.92, *Evc ω*=1.93, *Gli2 ω*=1.10; Supplementary Table 7). One of these three genes, *Gli2*, showed a statistically significant difference (*ω*=1.10 (Galapagos branch) vs. *ω*=0.11 (Background branch), P = 2.4×10-3; Supplementary Table 7). In contrast, *GU2* showed no signs of selection in the Galapagos Cormorants sister group (*P. auritus* and *P. brasilianus*) (*ω*=0.11 (Galapagos branch) vs. *ω*=1×10-4 (Background branch), P = 0.46). As a control, we also analyzed *Gli3*, the partially redundant paralog of *Gli2*, which also mediates Hh signaling but has no predicted function-altering variants in*P. harrisi* and found no evidence for positive selection (*ω*=0.04 (Galapagos branch) vs. *ω*=0.15 (Background branch), P = 0.11; Supplementary Table 7). These results suggest that positive selection could be partially responsible for the flightless phenotype of *P. harrisi*.

**Figure 4:**
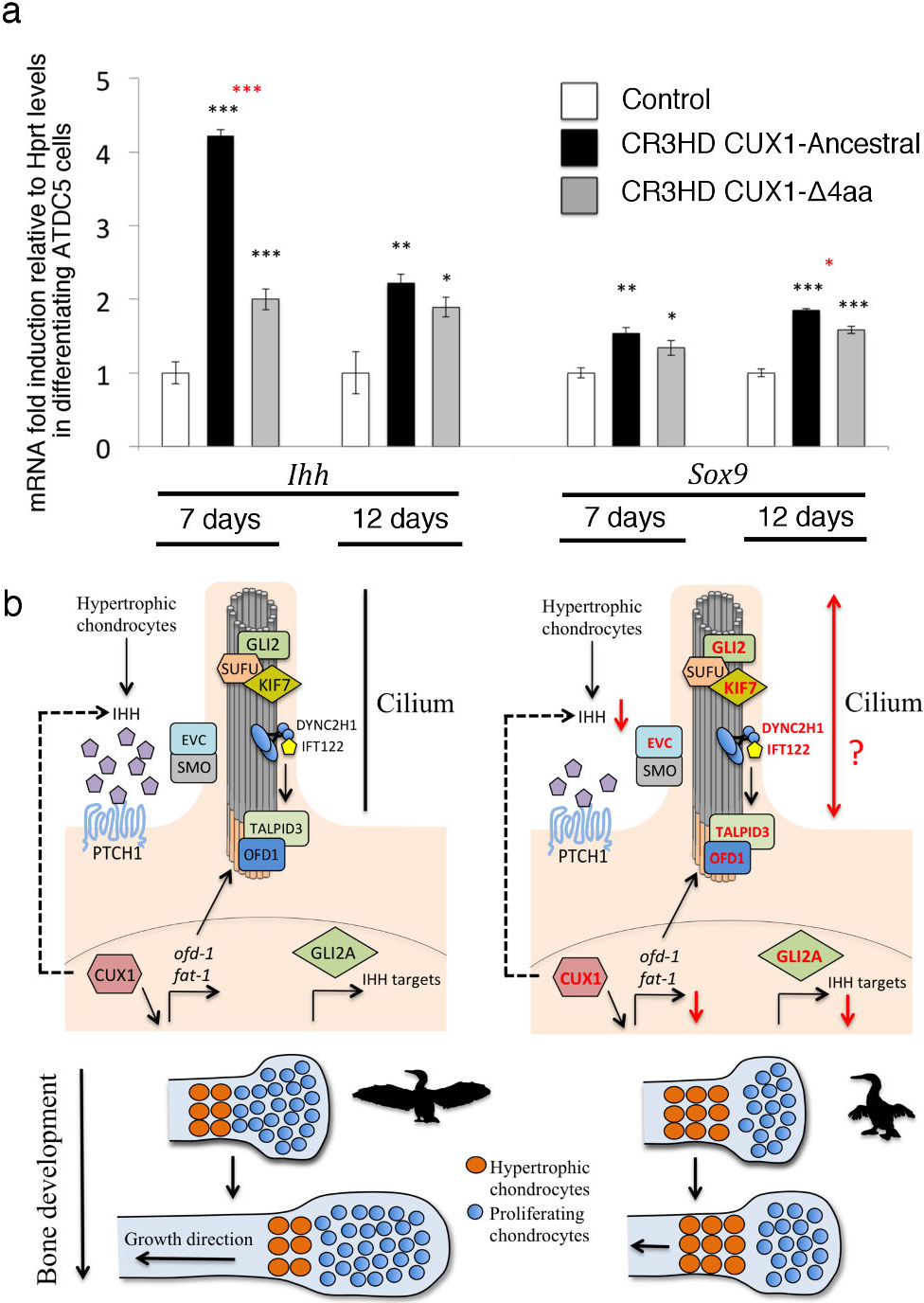
CR3HD CUX1 promotes chondrogenesis. a, ATDC5 control cells and cells carrying CR3HD CUX1-Ancestral or CUX1-Δ4aa variants were differentiated into chondrocytes. Gene expression levels were measured by RT-qPCR (n=4 biological replicates, each compromising 3 technical replicates) after 7 and 12 days. Error bars indicate standard errors. We used ANOVA and Tukeys HSD to test for statistical significance. Black stars indicate a significant difference between a CUX1 variant and the control. Red starts indicate a significant difference between CUX1-Ancestral and CUX1-Δ4aa variants. Absence of stars indicates no significant difference. (*) P<0.05, (**) P<0.01, and (***) P<0.001. b, Proposed mechanism for the reduction of wing size in*P. harrisi*. The left panel depicts the normal functioning of IHH signaling pathway in vertebrates. A selected number of proteins are shown for simplicity. OFD1 and TALPID3 are part of the basal body from which cilia are nucleated. textitOfd1 and talpid3 mutants are defective in cilia formation. In the presence of IHH secreted by hypertrophic chondrocytes, PTCH1 exits the cilium, where it cannot longer inhibit SMO. SMO is then activated and enriched in the cilium in association with EVC. KIF7 mediates the movement of GLI2/3-SUFU complex to the cilium. Then a series of molecular events leads GLI2/3 activation, transcription of IHH target genes, and chondrocyte proliferation. CUX1 promotes the expression of cilia related genes such as textitOfd1 and promoters chondrogenesis. The right panel depicts the state of the IHH pathway in *P. harrisi.* Proteins in red are affected by function-altering variants in *P. harrisi*. We predict these variants will affect both cilia formation and functioning leading to a reduction in IHH pathway activity. As a result, the pool of proliferating chondrocytes would decrease in wing bones and the number of hypertrophic chondrocytes would increase resulting in impaired bone growth.

## Discussion

The study of flightlessness evolution in the Rallidae family led Olson to propose in 1973 that flightlessness could be a fast-evolving heterochronic condition (Feduccia, 1999; Olson, 1973). Heterochrony, the relative change in the rate or timing of developmental events among species, is thought to be an important factor contributing to macroevolutionary change(Gould, 1977). Yet, virtually nothing is known about its genetic and molecular mechanisms. Diverse myological, osteological, and developmental observations strongly suggest that flightlessness in the Galapagos cormorant is caused by the retention into adulthood of juvenile characteristics affecting pectoral and forelimb development (a class of heterochrony known as paedomorphosis) (Livezey, 1992).

In this study we provide a genetic and molecular model that accounts for this heterochronic condition. We show that *Cux1*, a highly conserved transcription factor, has a four amino acid deletion in its regulatory domain. We molecularly dissected the consequences of this deletion and demonstrated that this variant impairs the ability of CUX1 to transcriptionally up-regulate cilia related genes (some of which are affected by variants themselves like *Ofd1* and *Fat1*) and promote chondrogenic differentiation. The wing and sternum of birds are some of the last structures to grow and ossify, and most of this growth occurs after hatching (Dunn, 1975; Olson, 1973). A premature arrest of bone growth due to reduced IHH signaling would mainly affect these organs. Thus, perturbations of cilia/Ihh signaling could be responsible for the reduction in growth of both wings and keel in the Galapagos cormorant. Of special interest is the gene *Fat1*, a target of Cux1 (Fig. 3d), which contains two function-altering variants (Table 1). *Fat1* −/- mouse mutants show selective defects in face, pectoral and shoulder muscles but not in in hindlimb muscles (Caruso et al., 2013). Thus, variants in *Fat1* could explain, in addition, the underdeveloped pectoral muscles of*P. harrisi*.

Although we have identified multiple variants that very likely contribute to the flightless phenotype of*P. harrisi*, we cannot exclude the contribution of other genes and pathways that our analysis may have missed, nor the contribution of non-coding regulatory variants (see Supplementary Note). Further characterization of the individual and joint contributions of the variants found in this study will help us to reconstruct the chain of events leading to flightlessness and to genetically dissect macroevolutionary change. We hypothesize that mutations in cilia or functionally related genes could be responsible for limb and other skeletal heterochronic transformations in birds and diverse organisms, including humans.

## Methods

**Library preparation and sequencing** DNA from*P. harrisi* was extracted from blood samples preserved in lysis buffer using the phenol-chloroform method. DNA from*P. auritus*., P. brasilianus and P. pelagicus was extracted from fresh tissues using the Qiagen DNA extraction kit and genomic tips (500/G). Before preparing DNA libraries, we checked the integrity of the samples by running 200 ng of DNA on a 0.6% agarose gel and observing a single high molecular weight band without smear. We generated libraries using three different protocols depending on the insert size. See Supplementary Table 2. For the smallest insert libraries (160bp) we used the Nextera protocol (Illumina). For mid-sized inserts (380, 430, and 670bp), we used the Truseq Nano protocol (Illumina). For the mate-pair libraries (2, 3, 4, 6, and 9 kb) we used the Nextera Mate-pair protocol (Illumina). We followed standard protocols with the following exceptions. For Nextera and Truseq protocols, we performed agarose size selection to guarantee a small variance in insert size. For Nextera Mate-pair protocols we used the Gel-plus protocol starting from 4 μg of genomic DNA. To shear circularized DNA we used 130μL microTube AFA Snap-cap vials and the Covaris M220 sonicator. All libraries were quantified using Qubit HS and the qPCR KAPA library quantification kit. Small insert and mate-pair libraries were sequenced using 100bp paired-end reads on an Illumina HiSeq 2000 at the UCLA Broad Stem Cell Center. The 170bp, 380bp insert, and mate-pair libraries were sequenced using a loading concentration of 12pM. 430bp insert and 670bp insert libraries were sequenced using 15pM and 18pM.

**Genome assembly** Sequence reads were filtered to meet genome assembly standards. We removed all duplicated reads. We used Trimmomatic (Bolger et al., 2014) to remove reads containing adapter sequences. We filtered out those reads having 40% of bases with Illumina Q-score 7 (Nextera and Truseq Nano libraries) and 60% of bases with Illumina Q-score 7 (Nextera mate-pair libraries). We used NextClip (v0.8)(Leggett et al., 2014) to remove the junction adapter from mate-pair libraries. We corrected errors (with the exception of mate-pair libraries) using SOAPec (v2.01), an error correction tool based on k-mer frequency. We assembled genomes using SOAPdenovo2 (Luo et al., 2012). We assembled the four genomes were using a k-mer size of 35. Small and mid-sized insert libraries were used for contig and scaffold assembly, and mate-pair libraries were used only for scaffolding. Once the assemblies were completed, we used GapCloser (v1.12) to fill the gaps (poly-N stretches) in the final assembly.

**Genome completeness evaluation** We ran the CEGMA pipeline (v2.4)(Parra et al., 2009) using: geneid (v1.4), genewise (v2.4.1), HMMER (v3.0) and NCBI Blast (v2.2.25+). This pipeline provides an unbiased estimate of assembly completeness based on its ability to retrieve a pre-defined subset of 248 highly conserved eukaryotic genes (CEGs). All genomes were analyzed with default parameters and the option-vrt (optimization for vertebrate genomes). The number of uniquely annotated proteins was calculated using the same pipeline used for the annotation of the genomes. To make a fair comparison between our genomes and those in the literature, we re-annotated 17 previously published bird genomes using exactly the same pipeline. The CEGMA score was highly correlated with the total number of uniquely annotated proteins (Fig. 2d).

**Homology-based annotation** We annotated the genomes using a homology-based approach from a set of 14,052 high confidence chicken proteins, for which there is evidence of expression in chicken and/or conservation in other species (Ensembl release 78, Galgal4). This allowed us to annotate 13,677 (97.3%), 13,652 (97.2%), 13,652 (97.2%), and 13,116 (93.3%) unique proteins in*P. harrisi,P. auritus,P. pelagians* and *P. brasilianus*, respectively. We also used this pipeline to predict orthologous proteins in the recently published Great cormorant (*Phalaaroaorax carbo*) genome (Jarvis et al., 2014) and found only 11,994 (85.0%) of them. The lower completeness of the P. carbo genome is most likely due to the lower coverage and lack of mate-pair libraries sequences. First, we used genBlastA (v1.38)(She et al., 2009) to define high-scoring segment pairs (HSPs) for each protein with options-c 0.3-e 0.00001-gff-pid-r 1. Then, we used exonerate (v2.2.0) (Slater and Birney, 2005) to build a gene model for every chicken protein using the genomic region defined by the HSP and 100kb of additional sequence surrounding it. We selected the model with the best score for each protein. If more than one model had the same score, we choose one at random. Our final datasets contain only one putative ortholog per chicken protein. A protein is defined as present in a given assembly if at least 30% of the length of the orthologous chicken protein can be mapped.

**Transcriptome-based annotation** We isolated the developing wings from *aP. auritus* embryo (Supplementary Fig. 5c). We estimated the embryo to be 15 days based on a photographic guide(Powell et al., 1998). This is the same individual whose DNA was used to generate the reference genome. Total RNA was extracted from the wings using the RNeasy Isolation kit (QIAGEN). RNA quality (RIN: 8.8) was determined using the RNA ScreenTape on an Agilent 2200 TapeStation (Agilent Technologies). We used the TruSeq Stranded mRNA kit (Illumina) to prepare the library and sequenced it using 100bp paired-end reads on an Illumina Hiseq 2000. We used 120 million paired-end reads (without removing duplicates) to de novo assemble the transcriptome using Trinity (v2.0140717) (Grabherr et al., 2011). We identified 63,500 candidate coding regions within 211,607 contigs using Transdecoder. 50.305 out of the 63,500 protein isoforms were further selected because they retrieved a blast hit against a database of the 14,052 high quality Chicken proteins (e =1×10-6). These 50.305 isoforms correspond to 11,753 of the 14,052 unique chicken proteins. We finally annotated the genome of*P. harrisi* and *P. pelagicus* (control for gene enrichment) using the 50.305 predicted protein isoforms from *P. auritus*. in an analogous way to the homology-based approach using genBlastA and exonerate.

**Phylogenetic tree and divergence estimates** We obtained common orthologs from the whole genome sequences of five different avian species: G. gallus (outgroup), N. nippon,*P. harrisi,P. auritus*, P. brasilianus, P. pelagicus, and P. carbo. We used BLAST to identify wrongly assigned orthologs and removed those genes from the dataset. Then, we aligned orthologs with MUSCLE(Edgar, 2004) and extracted all the fourfold degenerate sites. Our final dataset consisted of 9,641 orthologs containing a total of 1,717,714 fourfold degenerate sites. We used MrBayes (v3.2.5)(Ronquist et al., 2012) to reconstruct the phylogeny and estimate species divergence time using the following parameters: lset nst=6 rates=invgamma; prset brlenspr=clock:uniform, prset clockvarpr=igr, prset igrvarpr = exponential(2), prset clockratepr = normal(0.002,0.001), prset treeagepr = offsetexponential(0,500), calibrate tree = truncatednormal(70, 88,3). We used a GTR + I + Γ nucleotide substitution model. An estimate for substitution rate of fourfold degenerate sites in birds (0.002 substitutions per site per million years) was obtained from Green et al.(Green et al., 2014). We used the independent gamma rates (IGR) relaxed clock model. Due to the lack of an appropriate fossil for this phylogeny, the only calibration applied was the split between Galloanseres and Neoaves based on the estimate by Jarvis et al.(Jarvis et al., 2014). The analysis was run for 5 million generations.

**Heterozygosity calculation** For each species, we aligned using bwa (v0.6.2)(Li and Durbin, 2009) 20X of reads, originally used to generate the assembly. We then use samtools (v0.1.18)(Li et al., 2009) to call variants and generate vcf files. We used R to extract the number of heterozygous site. We restricted our counts to only single nucleotide polymorphisms (SNPs) without ambiguous bases, with a quality score >150, and depth < 100X. The heterozygosity in each genome is calculated as the number of heterozygous sites divided by total size of the assembly.

**Variant effect predictions** We ran PROVEAN (v1.1.5)(Choi et al., 2012) using: CD-HIT (v4.5.8), BLAST 2.2.30+ and NCBI nr database (downloaded February 2015). PROVEAN is based on evolutionary conservation and it has been validated with known human disease causing mutations and outperforms other prediction tools in experimental evolution studies that mimic the process of gradual accumulation of mutations encountered in nature(Rockah-Shmuel et al., 2015). PROVEAN needs two inputs: the reference protein sequence in fasta format and an amino acid variant file. In order to generate the variant file, the predicted reference and alternative proteins were first aligned using MUSCLE (v3.8.31) and then a custom Python script was used to call substitutions, deletions and insertions following the HGVS (Human Genome Variation Society) recommendations. We curated our predictions to minimize the presence of spurious variants. First, we re-blasted all the predicted proteins for each cormorant species against a database consisting of the initial 14,052 high quality chicken proteins. We removed those instances when a paralog was wrongly assigned as an ortholog of the chicken protein. Second, for each cormorant genome, we aligned 20X of the reads used to generate the assemblies back to the assemblies using bwa (v0.6.2). Then we called variants using samtools (v0.1.18). We searched for sites where 100% of the original reads supported a different nucleotide than the one in the final assembly. We assumed those SNPs corresponded to errors in the assembly. We removed genes that contained assembly errors within their coding regions from the final analyses.

**Gene enrichment analysis** We used the g:Profiler (Reimand et al., 2011) package for R. For the analysis of function-altering variants in *P. harrisi*. we set the list of 14,052 chicken proteins used to annotate the genomes as the background list. Our dataset consisted of three homology-based (*P. harrisi* vs. *P. auritus,P. harrisi* vs. *P. brasilianus*, and *P. harrisi* vs. *P. pelagicus*) and one transcriptome-based (*P. harrisi* vs. *P. auritus*) variant prediction datasets. We defined a set of proteins containing function-altering variants (PROVEAN score < −5) in at least one of our four prediction datasets (1129 proteins of which 1101 have a known ontology). This strategy was conservative and guaranteed that variants derived exclusively from the transcriptome prediction were included in the analysis. The enrichment for human disease categories was derived from annotations from the Human Phenotype Ontology (HPO), a standardized vocabulary of phenotypic abnormalities encountered in human disease. To check for the specificity of enrichments for syndromes affecting limb development in*P. harrisi*, we performed gene enrichment analysis in the union set of variants predicted to be function-altering in *P. auritus*, P. brasilianus and P. pelagicus compared to *P. harrisi* (PROVEAN score > 5) in our four prediction datasets. Categories on this set did not show any limb related syndrome or overlap with the previous categories, (Supplementary Table 4b). As a second specificity control, we repeated our prediction pipeline, but instead of*P. harrisi*, we used P. pelagicus as a reference species. Enriched human disease categories in P. pelagicus did not show any overlap with the ones from*P. harrisi*. (Supplementary Table 4c). For the enrichment analysis of CUX1 target genes in Hs578t cells (breast cancer cell line), we used the list of all genes contained in the array probes as the background. We applied the g:SCS method for estimating significance and correcting for multiple testing.

**Confirmation of function-altering variants** We performed Sanger sequencing to confirm the presence of predicted function-altering variants in*P. harrisi* and its absence in the three closely related cormorants. A list of the primers used can be found in Supplementary Table 10. To determine if a variant was fixed in the Galapagos Cormorant population, we genotyped by Sanger sequencing another 20 different individuals from two of the most divergent populations: Cabo Hammond (10) and Caones Sur (10) (Duffie et al., 2009). Given that the total population of the Galapagos Cormorant is 1,500 individuals, we considered that if 20/20 genotyped individuals were homozygous, then the variant was most likely fixed in the population.

**Electrophoretic Mobility Shift Assay (EMSA)** We digested the plasmid pMAL-c5X-His (NEB) with NcoI and SbfI-restriction enzymes and performed Gibson assembly to insert gBlocks (IDT) coding for E. coli codon optimized CR3HD CUX1-Ancestral or CUX1-Δ4aa variants. Expression of this plasmid results in a fusion protein containing the Maltose binding protein (MBP) on the N-terminal to increase solubility, the CR3HD domain, and a 5X His tag on the C-terminal for protein purification. Proteins were expressed in E. coli and purified at the Protein Expression Technology Center, UCLA. After protein elution, we performed MWCO 50 kDa dialysis using the Pur-A-Lyzer Maxi 50000 kit (Sigma) and 1X Binding Buffer. Purified protein was quantified using the Bradford Protein Assay Kit (Thermo-Scientific) and kept at −80C. Expected size and protein purity was confirmed running 10 ng on a 10% Bis-Tris polyacrylamide gel under denaturing conditions and performing silver staining with Pierce Silver Stain kit (ThermoFisher). In addition, purified proteins were recognized by MBP (NEB) and (M-222) CUX1 antibodies, as expected (not shown). To perform gel-shifts we used the LightShift Chemiluminescent EMSA kit from Thermo Scientific. The experimental conditions for CR3HD CUX1 binding were adapted from a previous study(Moon et al., 2000). Purified CR3HD CUX1-Ancestral or CUX1-Δ4aa protein was pre-incubated for 5 minutes with 1μg of poly(dI-dC) and 3.3μg of BSA in 25mM NaCl, 10mM Tris, pH 7.5, 5nM EDTA, 5% Glycerol, 1mM DTT, 1mM MgCl2. 20fmol of Biotin end-labeled target dsDNA (TCGAGACGGTATCGATAAGCTTCTTTTC) were added to the mix and incubated at RT for 20 minutes.

**Luciferase assay** We adapted a previously validated chloramphenicol acetyltransferase (CAT) reporter assay to test the repressor activity of the C-terminal tail of CUX1(Mailly et al., 1996). We generated a luciferase reporter plasmid by cloning a gBlock (IDT) containing 7xUAS binding elements followed by the thymidine kinase (tk) promoter using Gibson assembly into the pGL4.15[luc2P/Hygro] plasmid (Promega) previously digested with XhoI and HindIII restriction enzymes. We then generated GAL4DBD protein fusion plasmids following two steps. First, we cloned a gBlock (IDT) coding for a human codon optimized GAL4 DBD (1:147 aa) into a pcDNA3.1(+) (Thermo-Fisher) mammalian vector previously digested with HindIII and EcoRI using Gibson assembly. We then used this plasmid as a backbone to introduce the CUX1 C-terminal domains containing the repressor activity. We then digested the previously modified a pcDNA3.1 plasmid with XhoI and XbaI, and used gBlocks (IDT) to clone the CUX1 C-terminal domain Ancestral or Δ4aa variants using Gibson assembly. We transfected COS-7 cells growing in 12 well plates under standard conditions with Lipofectamine 3000 (ThermoFisher) using 600ng of GAL4DBD:CUX1-Cterm plasmid, 200ng of 7xUAS/tk:luc2 reporter, and 2ng of pGL4.75[h/Rluc/CMV] as a normalization control. We measured reporter activity 48 hours after transfection. All transfections and measurements were performed in triplicate.

**Western blot** We incubated commercial mallard eggs (Metzer Farms, CA) and extracted the wings from developing mallard embryo after 22 days of incubation at 37.5C and 55% relative humidity. We incubated the tissue in RIPA lysis buffer supplemented with Halt Protease Inhibitor Cocktail (Thermo Scientific). The lysate was further subjected to mechanical lysis and sonication at 4C. Protein concentration was determined using the Bradford Protein Assay Kit (Thermo-Scientific). 20μg of protein were loaded into a 10% Bis-Tris polyacrylamide gel under denaturing conditions. Sample was transferred to a PVDF membrane and blocked for 2 hours in 5% BSA PBST buffer at RT. The membrane was incubated overnight at 4C with 5% BSA PBST and a 1:1000 dilution of rabbit polyclonal M-222 CUX1 antibody (200 μg/mL) (Santa Cruz Biotechnology). After 3 washes with PBST buffer, the membrane was incubated with a 1:10,000 dilution of Peroxidase-conjugated Goat Anti-Rabbit IgG (H+L) (Jackson ImmunoResearch) in PBST for 1h at RT.

**ATDC5 cell line generation and differentiation** CR3HD CUX1-Ancestral and CR3HD CUX1-Δ4aa mouse codon optimized genes were cloned into a lentiviral vector to produce N-terminal HIS-tag fusion proteins under a CMV promoter. ATDC5 cells were transduced with lentiviral particles and stable cell lines and were generated using puromycin selection. A control ATDC5 line was generated under identical conditions but using lentiviruses carrying an empty vector. Stables cell lines were generated by Cellomics Technology, MD, USA. Immunostaining against the HIS tag revealed that both variants have strong nuclear localization, indicating that the Galapagos Cormorant variant does not affect the subcellular localization of CUX. Cells were plated at 1.5105 cells/well in 24-well plates, and were cultured using DMEM+10% FBS and 3 μg/mL puromycin. To differentiate cells we cultured cells in chondrogenic differentiation medium:-MEM containing 5%FBS, 200 μg/ml ascorbic acid, 60 nm Na2SeO3, 10 μg/ml transferrin, 1% antibiotic. After 7 and 12 days in culture, cells were lysed and total RNA was isolated using the RNeasy Isolation kit (QIAGEN).

**Real time quantitative PCR** RNA from ATDC5 cell lines was extracted using the RNeasy Isolation kit (QIAGEN). cDNA was prepared using Superscript III Reverse Transcriptase (ThermoFisher) using Poly-T primers. All gene specific primers were designed to span exon-exon junctions and/or exon-intron boundaries when possible. A list of the primers used can be found in Supplementary Table 11. We validated primers by constructing standard curves and checking for amplification efficiency. All measurements were performed in at least three biological replicates per condition. As a reference gene, we used Hprt, which was previously validated in ATDC5 cells (Zhai et al., 2013). Importantly, Hprt levels were not affected in Hs578t cells overexpressing CUX1 nor expressing CUX1 shRNA and have been previously used to validate Cux1 target genes. The use of another validated reference gene, Ppia, gave identical results to Hprt. All measurements were performed using the SYBR Green I master mix (Roche) in a LightCycler 480 Real-time PCR System (Roche)

**Statistical test for positive selection** Ortholog coding sequences were obtained from our assemblies and the NCBI database. Nucleotides sequences were aligned using protein alignments as a reference with PAL2NAL (Suyama et al., 2006). Alignments were visually inspected and taxa containing errors were discarded if low quality. We estimated the ratio of non-synonymous to synonymous substitutions (*ω* = dN/dS) using a maximum likelihood framework and a branch model of adaptive evolution implemented in CODEML (PAML v4.8) (Yang, 2007). We used the likelihood ratio test to compare a null model (M0) that estimates an average for a given gene across all branches in the phylogeny and an alternative two-ratio branch model that estimates one for the branch of interest and another for the background branches. Our phylogeny consisted of 36 taxa: 28 birds (*Columba livia, Acanthisitta chloris, Manacus vitellinus, Corvus cornix, Taeniopygia guttata, Nestor notabilis, Calypte anna, Meleagris gallopavo, Gallus gallus, Cuculus canorus, Mesitornis unicolor, Struthio camelus, Anas platyrhynchos, Falco peregrinus, Falco cherrug, Egretta garzetta, Nipponia nippon, Charadrius vociferous, Phalacrocorax pelagicus, Phalacrocorax carbo, Phalacrocorax auritus, Phalacrocorax harrisi, Phalacrocorax brasilianus, Fulmarus glacialis, Aquila chrysaetos, Haliaeetus leucocephalus, Aptenodytes forsteri, and Pygoscelis adeliae*), 6 other reptiles (*Python bivittatus, Anolis carolinensis, Gekko japonicus, Chelonia mydas, Alligator sinensis, Alligator mississippiensis*) and 2 mammals (*Homo sapiens and Mus musculus*).

## Author contributions

A.B. and L.K. conceived the study. A.B. coordinated the collection of samples, prepared libraries, assembled and annotated genomes, and performed all the analyses and experiments. P.W., A.R., C.V., and P.P. provided DNA or tissue samples. W.W. and A.B. carried out ATDC5 cell line experiments supervised by K.L. AB. and L.K. wrote the manuscript. All authors discussed and agreed on the final version of the manuscript.

## Acknowledgement

A.B. is funded by the Jane Coffin Childs Memorial Fund for Medical Research. L.K. is funded by the Howard Hughes Medical Institute (HHMI). A.R. is funded by the U.S. Geological Survey through the Wildlife Program of the Ecosystems Mission area. Any use of trade names is for descriptive purposes only and does not imply endorsement by the U.S. Government. C.V. is funded by FONDECYT (11130305). We thank Christopher Koo and Dr. Mark Arbing at the UCLA-DOE Protein Expression Laboratory Core Facility for protein purification and Dr. Suhua Feng at the Broad Stem Cell Research Center High Throughput Sequencing Core for assistance. We thank Dr. Carlos Valle (USFQ) for advice. We thank Dr. Kimball Garrett (Natural History Museum of Los Angeles), Dr. Kevin Burns (San Diego Natural History Museum and SDSU Museum of Biodiversity), and Dr. Rebecca Duerr (International Bird Rescue) for providing samples used in preliminary stages of this study.

## Supplementary Note

### Analysis of the 23 proteins with the strongest function-altering variants in the Galapagos Cormorant

We curated the list of 23 proteins (22 unique genes; the gene ankyrin repeat domain-containing protein 27 is represented by two isoforms) predicted to have the strongest function-altering variants in *P. harrisi* across datasets (PROVEAN score < 12.5). First, we curated the annotation of each of these proteins. In 8 cases out of the 23 proteins, the predicted variant could be satisfactory explained by an error produced during the annotation process. These cases usually involved highly repetitive and poorly conserved domains that resulted in large deletions (Extended Data Table 2). In 2 additional cases, the error was due to an assembly issue because exons predicted to be missing were actually present in another scaffold or contig. We verified the presence of the putative missing exons for ADNP-homeobox-2 and R-spondin-2 using Sanger sequencing. 13 proteins (12 unique genes) had variants that could not be explained by assembly nor annotation artifacts. We designed primers for all of them. We successfully obtained PCR products spanning the variant in question for 9 proteins (8 unique genes). We confirmed variants 7 out of the 8 genes by Sanger sequencing. The variant in FAM210A was a false positive. 2 out of the 7 genes with confirmed variants were discussed in the main manuscript (Cux1 and Lgals3). The remaining 5 genes were: Torsin family 1 member B-like precursor, Ankyrin repeat domain-containing protein 27, Angiomotin like 2, MICAL-like protein, and NEDD4 binding protein 2. Of these 5 genes, only Angiomotin like 2 (Amotl2) has known role in development. Loss of Amotl2 in zebrafish leads to over-proliferation of the lateral line primordium (Agarwala et al. 2015). This effect is thought to be mediated by the Hippo and Wnt/β-catenin pathways. Interestingly, Fat and Dachsous, which are both mutated in *P. harrisi*, are the upstream activators of the Hippo pathway in Drosophila, in addition to their role in PCP. Although it is tempting to speculate about the contribution of Amotl2 to the flightless phenotype of *P. harrisi*, more experimental evidence needs to be gathered. Variants in 4 genes could not be amplified by PCR in any cormorant despite multiple attempts and trying different concentrations of DMSO and several Tm. These genes were characterized by a high GC content (in 3 cases > 65% GC). A high GC content is not only problematic for PCR, but also likely to affect the assembly of these genes from Illumina short reads. Thus, the high GC content of some genes may cause the calling of spurious variants. Despite the lack of confirmation of these variants, none of these 4 proteins has a known role in limb development or morphogenesis (Extended Data Table 2).

### Analysis of genes that could not be annotated in the Galapagos Cormorant

375 out of the 14,052 high quality chicken proteins (2.7%) did not pass the detection threshold (>30% coverage of query protein length) when annotating the genome of *P. harrisi.* The failure to detect these proteins could be due to both technical and biological factors. For example, incorrect or incomplete assembly of regions of the genome could result in different parts of a gene mapping to diverse scaffolds and/or contigs. If any of the individual scaffolds or contigs contains less than 30% of the query protein, then our pipeline would miss that gene. This technical issue is more likely to affect large genes and genes within repetitive regions that are hard to assemble. In addition, it is possible that the genes not detected in the Galapagos Cormorant were lost in the cormorant lineage or are present only in the chicken lineage. Moreover, some chicken genes may be incorrectly annotated causing further issues. To gain insights into the nature of the “missing” genes in *P. harrisi*, we first asked how many of the 375 proteins not annotated in *P. harrisi* were also missing in the other three cormorant genomes (*P. auritus, P. brasilianus* and *P. pelagicus*). The large majority of the genes not annotated in *P. harrisi* (84%, 316 out of 375) did not, in turn, pass the annotation threshold in the other three sequenced cormorants. Moreover, 43.1% of proteins missing in all four cormorants were annotated as “Uncharacterized proteins” in contrast to 5.2% in the original data set. These results suggest that a large fraction of the chicken proteins not annotated in *P. harrisi* and the other cormorants are specific to chickens or erroneous gene models. 59 proteins could not be annotated in *P. harrisi* but were annotated in at least one other cormorant genome. BLAST search in of the 59 chicken proteins *P. harrisi* retrieved multiple high scoring hits in different scaffolds/contigs per protein, suggests that the large majority of “missing” genes in *P. harrisi* are simply not correctly assembled. Moreover, only 17 out of these 59 proteins were annotated in all three flighted cormorants, but were missing from *P. harrisi* (Supplementary Table 8). None of 17 the putative “missing” proteins are a likely candidate to influence wing development based on gene ontology, known expression patterns and available mutant phenotypes. However, we cannot fully discard their contribution.

### Analysis of genes with putative premature stop codons in the Galapagos Cormorant

Premature stop codons are considered to be highly detrimental because they can lead to truncated proteins or transcript degradation mediated by Nonsense-mediated mRNA decay (NMD). Genes affected by a premature stop codon in the Galapagos Cormorant thus deserve special attention. We annotated Galapagos Cormorants proteins based on the presence or absence of premature stop codons relative to the orthologous chicken stop codon. We found that 939 out of 13,652 of *P. harrisi* proteins (6.9%) were affected by a putative premature stop codon. The large majority of these stop codons, 867 out of 939 (92.3%), were shared with at least one other cormorant. Only 72 proteins had a predicted premature stop codon in *P. harrisi* that was not present in any of the other cormorants’ orthologs. We manually curated the annotation of these 72 proteins comparing the annotations of *P. harrisi* to that of its flighted cormorant relatives. The majority of these premature stop codons (69%, 50 out of 72) were found to be artifacts caused by issues with the genome assembly and/or annotation. These erroneous stop codons were largely located close to missannotated exons and introns. The remaining 21 proteins (18 unique genes) could indeed contain a premature stop codon. Among these 21 proteins we found a putative premature stop codon in the gene CILP (cartilage intermediate layer protein), which was previously associated with osteoarthrosis. We Sanger sequenced the putative CILP premature stop codon, but it was a false positive. None of the other 20 proteins (17 genes) had a known role in neither limb development nor morphogenesis. (Supplementary Table 9).

### Analysis of ultraconserved noncoding elements (UCNE)

To evaluate the contribution of non-coding variants to the flightless phenotype of *P. harrisi*, we searched for chicken ultraconserved noncoding elements (UCNE) in the genomes of *P. harrisi* and its close relative *P. auritus*. UCNEs are enriched for regulatory elements and often act as transcriptional enhancers. Of the 4,351 chicken UCNEs, we identified 4,348 unique UCNEs in the genomes of both *P. harrisi* and *P. auritus* using BLAST. The 179 missing UCNEs were common to both genomes, therefore not a single UCNE was missing exclusively in *P. harrisi* but present in its flighted close relative. The divergence of no single UNCE was higher than 3% (percent identity) between *P. harrisi* and *P. auritus*, suggesting no major deletions or rearrangements within UCNEs.

**Figure S1:**
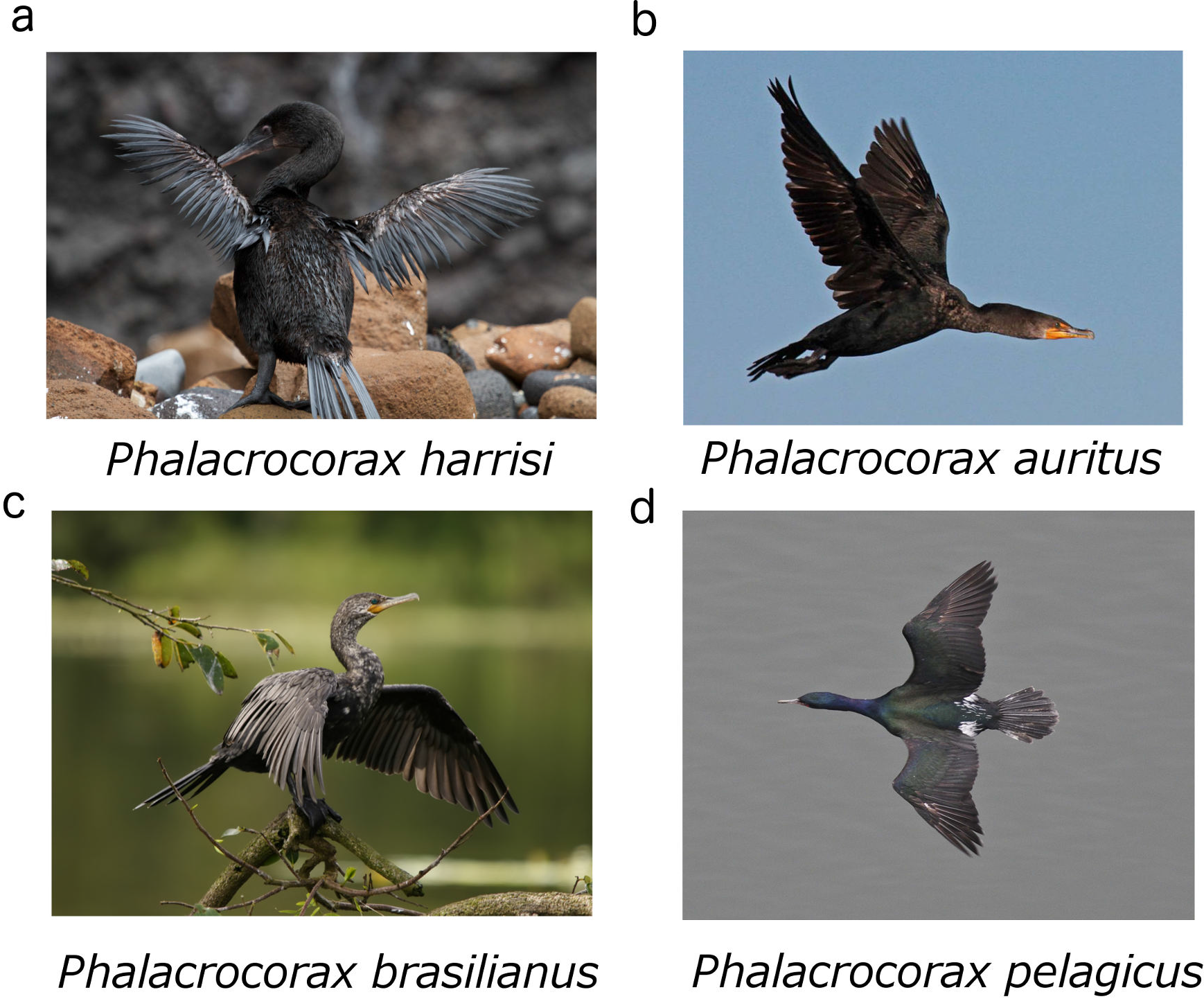
Sequenced cormorant species. a, Galapagos cormorant (Photo credit:Brian Gratwicke, flickr, CC BY 2.0) b, Double-crested cormorant (Photo credit: Dick Daniels, Wikipedia, CC BY-SA 3.0) c, Neotropical cormorant (Photo credit: Hans Hillewaert, Wikipedia, CC-BY-SA 4.0) d, Pelagic cormorant (Photo credit: U.S. Fish and Wildlife Service Headquarters, flickr, CC BY 2.0)

**Figure S2:**
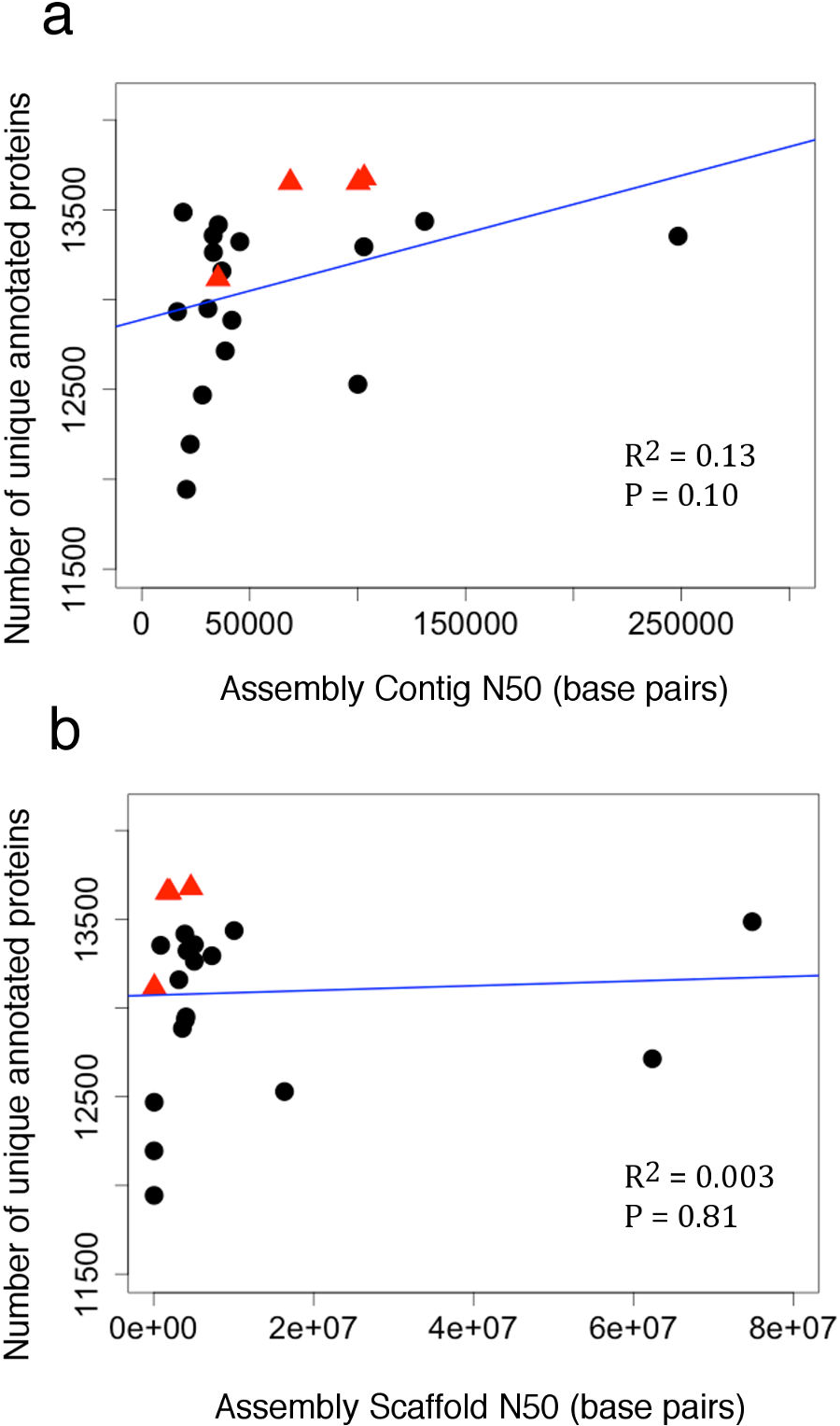
Contig and scaffold N50 are poor predictors of genome completeness. a, Neither contig N50 nor b, scaffold N50 predicts the total number of unique proteins in a given genome assembly. We used 14,052 high quality chicken proteins to annotate avian genomes. Blue line is the linear regression model. Genomes reported in this study are red triangles and other published avian genomes are black circles (see Supplementary Table 2 for full details).

**Figure S3:**
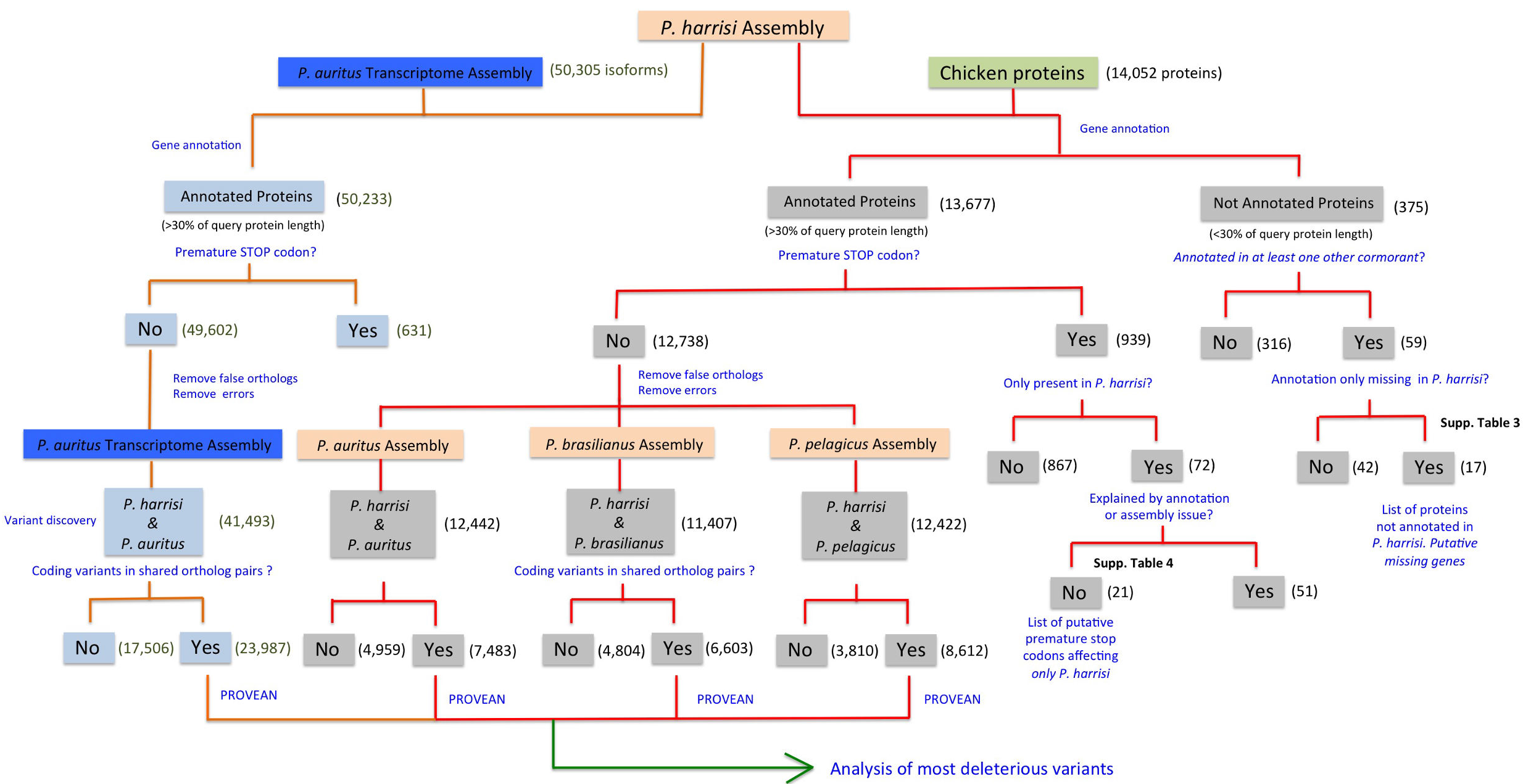
Flowchart for the discovery and characterization of coding variants in P. harrisi

**Figure S4:**
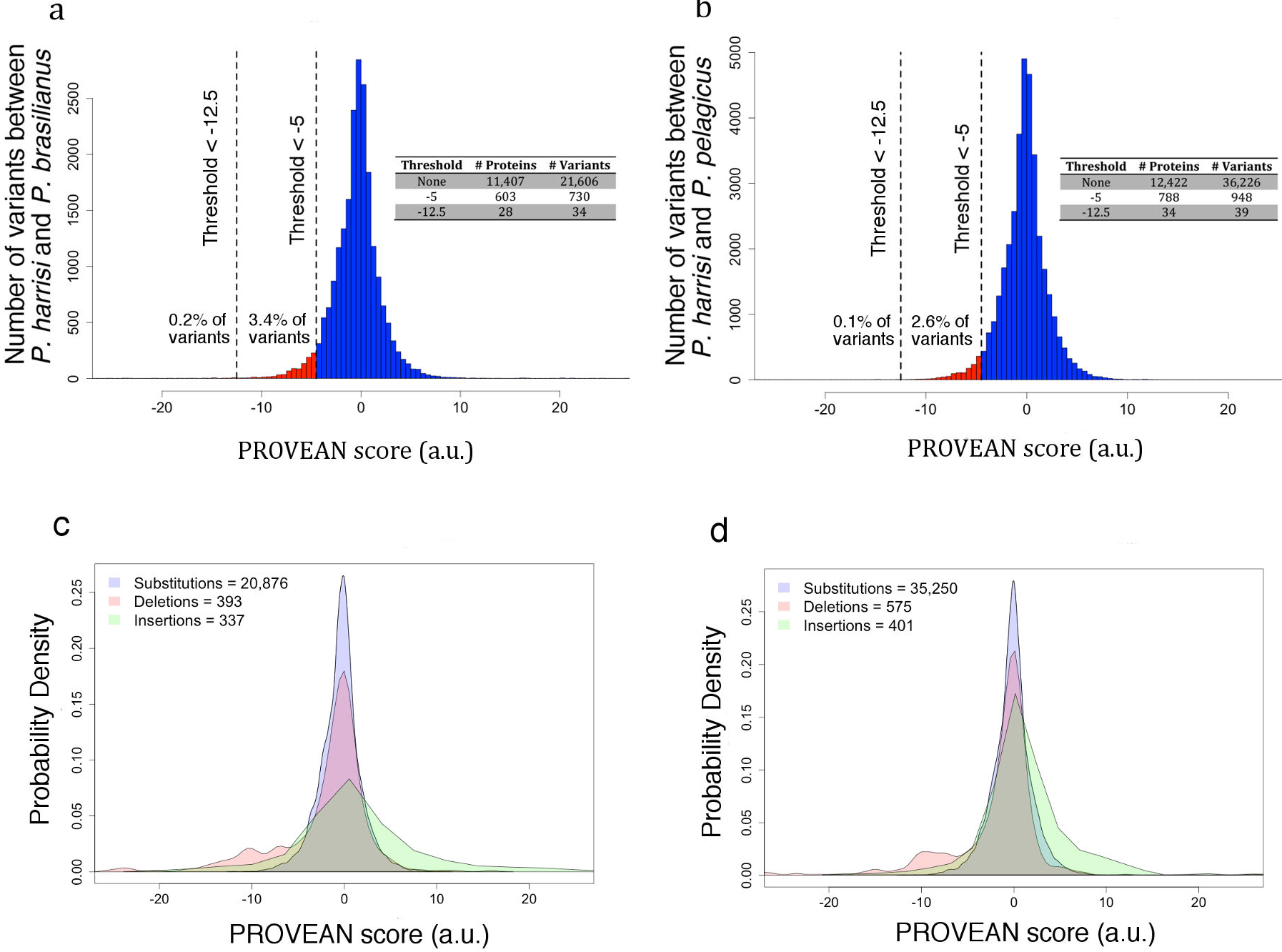
Distribution of the effect of variants between P. brasilianus and P. harrisi, and between P. pelagicus and P. harrisi. a, We used PROVEAN to predict the effect on protein function of 21,606 variants contained in 11,407 orthologous pairs between P. brasilianus and P. harrisi. 4,804 pairs contained no variants. b, Analogous to a, we used PROVEAN to predict the effect on protein function of 36,226 variants contained in 12,422 orthologous pairs between P. pelagicus and P. harrisi. 3,810 pairs contained no variants. PROVEAN score thresholds used in this study are drawn as vertical dashed lines. Number of proteins and variants found for each threshold are presented in inset table. c and d, Density of PROVEAN scores for each class of variant. The same variants presented in (a) and (b) were classified as single amino acid substitutions, deletions, and insertions. Number of variants in each class is indicated in the legend.

**Figure S5:**
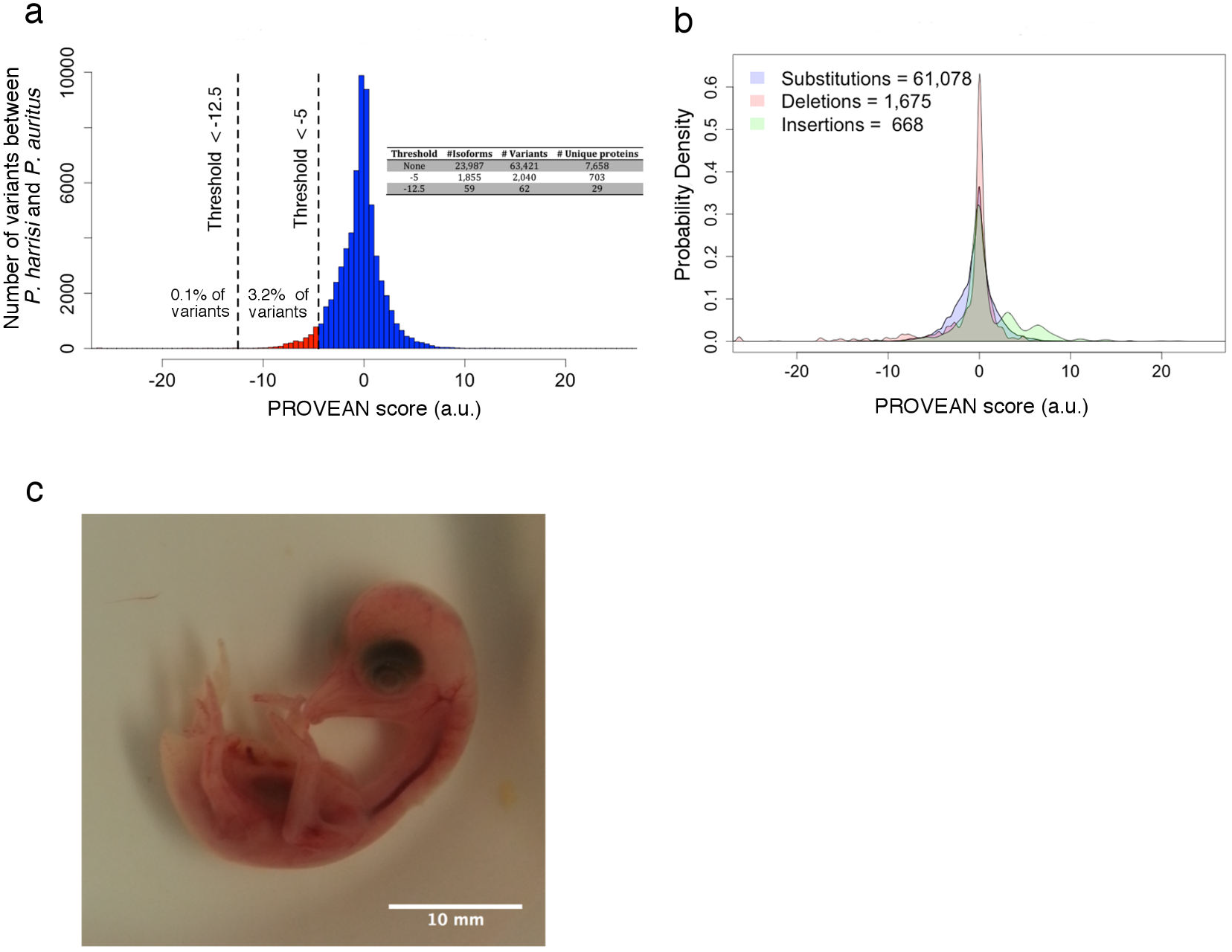
Distribution of the effect of variants between P. auritus and P. harrisi from transcriptome predictions. a, We used PROVEAN to predict the effect on protein function of 63,421 variants contained in 23,987 orthologous isoforms (7,658 unique proteins) between P. auritus and P. harrisi. These isoforms were predicted from a de novo transcriptome assembly from the wing of a developing embryo. PROVEAN score thresholds used in this study are drawn as vertical dashed lines. Number of proteins and variants found for each threshold are presented in inset table. b, Density of PROVEAN scores for each class of variant. The same variants presented in (a) were classified as single amino acid substitutions, deletions, and insertions. Number of variants in each class is indicated in the legend. c, The double-crested cormorant embryo used to generate the reference genome and wing transcriptome.

**Figure S6:**
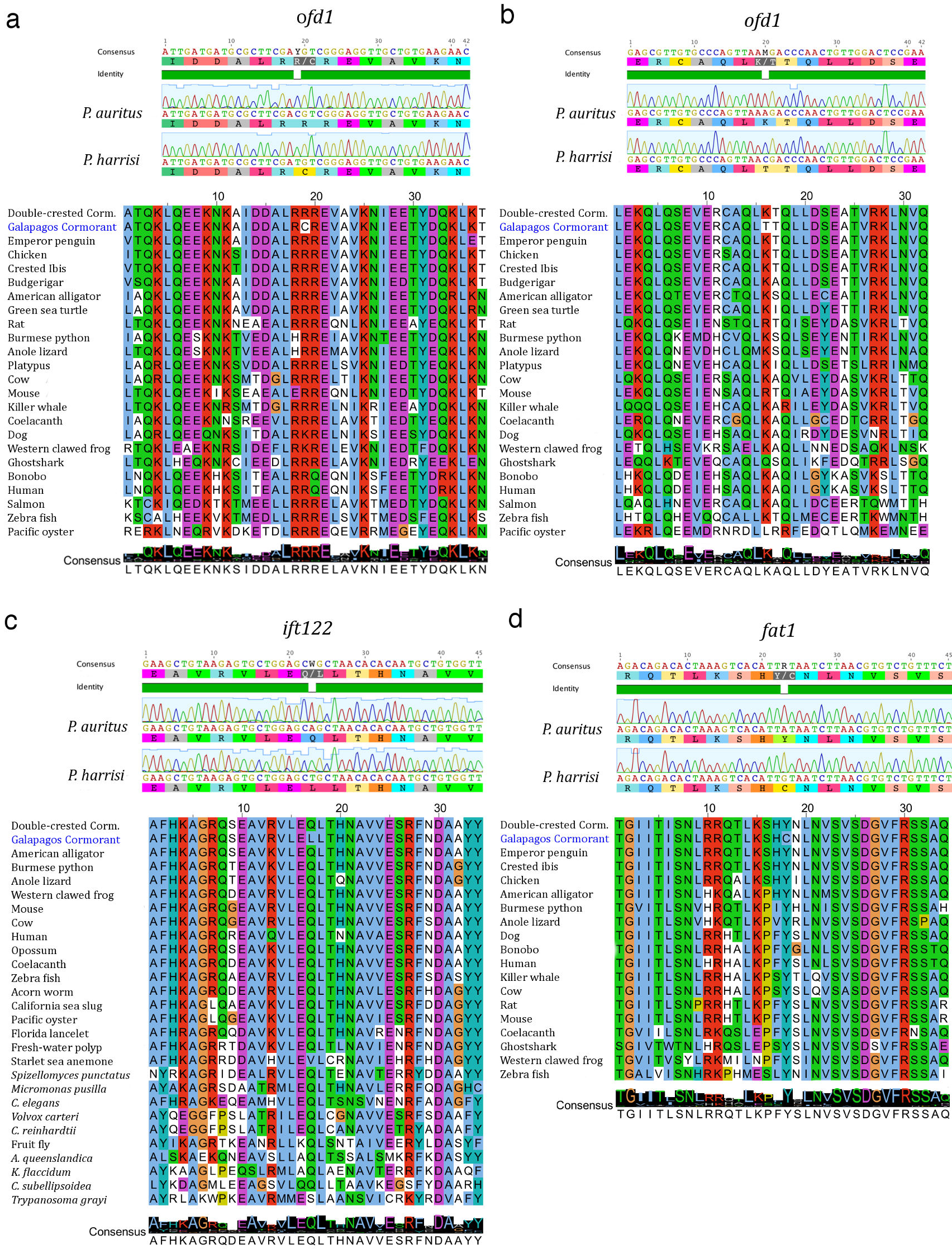
Sanger sequencing confirmation and evolutionary conservation of function-altering variants in P. harrisi. Sanger sequence confirmation and protein alignment for the following variants: a, OFD1 (R325C) b, OFD1 (K517T) c, IFT122 (Q691L) and d, FAT1 (Y2462C).

**Figure S7:**
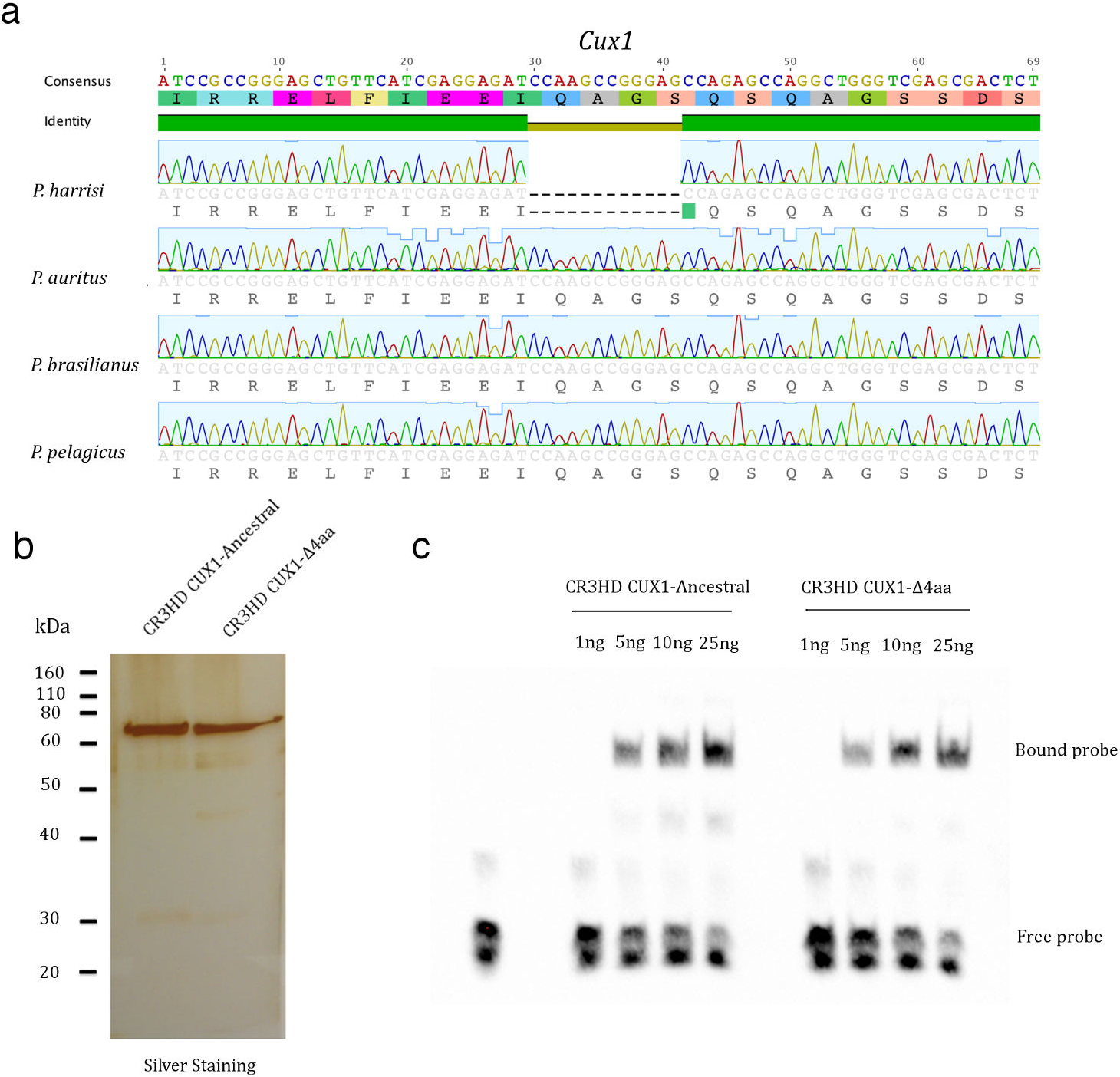
Cux1 deletion variant confirmation and CUX1 DNA binding experiments. a, We Sanger sequenced the area surrounding the predicted Cux1 deletion in P. harrisi, P. auritus, P. brasilianus and P. pelagicus. Alignment of the resulting sequences confirmed the 12bp deletion in P. harrisi. The deletion was not detected in the other cormorants. b, Silver staining of the purified CR3HD CUX1-Ancestral and CUX1-Δ4aa variants. The proteins matched the expected sizes for MBP-CR3HD CUX1-5xHisTag protein fusions. c, Electrophoretic Mobility Shift Assay (EMSA). Purified CR3HD CUX1-Ancestral and CUX1-Δ4aa variants were equally able to shift a labeled probed containing a CUX1 CR3HD binding motif.

**Supplementary Table 1.** Sequenced species and genome assembly statistics.

| Species | Common name | Location | Source of DNA | Sample Provider |
| --- | --- | --- | --- | --- |
| *Phalacrocorax harrisi* | Galapagos Cormorant | Punta Espinosa, Fernandina Island, Galapagos, Ecuador | Blood sample | Patricia Parker |
| *Phalacrocorax auritus* | Double-crested Cormorant | Minnesota Lake, Minnesota, USA<br>43.835 N,<br>-93.864 W | Embryo | Paul Wolf |
| *Phalacrocorax brasilianus* | Neotropical Cormorant | Naguilan, Valdivia, Chile.<br>-39.949759 S,<br>-73.303114 W | Fresh tissue | Claudio Verdugo |
| *Phalacrocorax pelagicus* | Pelagic Cormorant | Middleton Island, Alaska, USA<br>59.4375 N,<br>146.3275 W | Embryo | Andrew Ramey |

b. Assembly statistics
| Species | Assembly length (bp) | Contigs | Contig N50 (bp) | Scaffolds | Scaffold N50 (bp) |
| --- | --- | --- | --- | --- | --- |
| *Phalacrocorax harrisi* | 1,206,855,197 | 54,323 | 103,013 | 26,320 | 4,648,443 |
| *Phalacrocorax auritus* | 1,257,857,018 | 123,589 | 100,253 | 96,163 | 1,982,658 |
| *Phalacrocorax brasilianus* | 1,388,976,773 | 601,596 | 35,288 | 567,434 | 81,228 |
| *Phalacrocorax pelagicus* | 1,219,010,268 | 118,438 | 68,796 | 78,898 | 1,694,410 |

**Supplementary Table 2.** Sequenced libraries used for genome assembly.

**a, Libraries used for the assembly of the genome of *P. auritus*.** **b, Libraries used for the assembly of the genome of *P. brasiliensis*.**
| **Library** | **Protocol** | **Insert size**<br>**(bp)** | **Read**<br>**Length** | **Number of**<br>**raw reads** | **Number of**<br>**processed reads** | **Read**<br>**Coverage** | **Estimated Insert size**<br>**+/- sd (bp)** |
| --- | --- | --- | --- | --- | --- | --- | --- |
| **Pbrasil**<br>**300bp_V1** | Nextera | 160 | PE100 | 228101388 | 171723149 | 24.73 | 151 +/- 18 |
| **Pbrasil**<br>**500bp_V1** | Truseq<br>nano | 380 | PE100 | 168415774 | 158844156 | 22.87 | 366 +/- 29 |
| **Pbrasil**<br>**600bp_V1** | Truseq<br>nano | 480 | PE100 | 147789445 | 138608696 | 19.96 | 424 +/- 63 |
| **Pbrasil**<br>**850bp_V1** | Truseq<br>nano | 730 | PE100 | 194098352 | 187129176 | 26.94 | 610 +/- 133 |
| **Pbrasil**<br>**3kb_V1** | Nextera<br>Mate-pair | 3000 | PE100 | 126945773 | 60534605 | 8.72 | 2722 +/- 333 |
| **Pbrasil**<br>**4kb_V1** | Nextera<br>Mate-pair | 4000 | PE100 | 167215728 | 56881629 | 8.19 | 3689 +/- 558 |
| **Pbrasil**<br>**6kb_V1** | Nextera<br>Mate-pair | 6000 | PE100 | 156187926 | 42893892 | 6.18 | 5253 +/- 1052 |
| **Total coverage** |  |  |  |  |  | **170.99X** |  |

**c, Libraries used for the assembly of the genome of *P. harrisi*.**
| **Library** | **Protocol** | **Insert size (bp)** | **Read Length** | **Number of raw reads** | **Number of processed reads** | **Read Coverage** | **Estimated Insert size +/- sd (bp)** |
| --- | --- | --- | --- | --- | --- | --- | --- |
| **Pharrisi 300bp_V1** | Nextera | 160 | PE100 | 194488597 | 161153950 | 26.71 | 139 +/- 16 |
| **Pharrisi 500bp_V1** | Truseq nano | 380 | PE100 | 206270105 | 189615839 | 31.42 | 337 +/- 31 |
| **Pharrisi 600bp_V1** | Truseq nano | 480 | PE100 | 203159496 | 188494910 | 31.24 | 406 +/- 56 |
| **Pharrisi 850bp_V1** | Truseq nano | 730 | PE100 | 171622105 | 162525787 | 26.93 | 650 +/- 159 |
| **Pharrisi 2kb_V1** | Nextera Mate-pair | 2000 | PE100 | 149734456 | 79875422 | 13.24 | 1693 +/- 227 |
| **Pharrisi 2kb_V2** | Nextera Mate-pair | 2000 | PE100 | 188633347 | 50246924 | 8.33 | 1814 +/- 217 |
| **Pharrisi 3kb_V1** | Nextera Mate-pair | 3000 | PE100 | 178696882 | 82358300 | 13.65 | 2791 +/- 289 |
| **Pharrisi 4kb_V1** | Nextera Mate-pair | 4000 | PE100 | 210556792 | 102525061 | 16.99 | 3866 +/- 491 |
| **Pharrisi 4kb_V2** | Nextera Mate-pair | 4000 | PE100 | 181794395 | 40878684 | 6.77 | 3835 +/- 506 |
| **Pharrisi 6kb_V1** | Nextera Mate-pair | 6000 | PE100 | 192352397 | 46604124 | 7.72 | 5425 +/- 1017 |
| **Pharrisi 6kb_V2** | Nextera Mate-pair | 6000 | PE100 | 203957935 | 46683753 | 7.74 | 5345 +/- 969 |
| **Pharrisi 9kb_V1** | Nextera Mate-pair | 9000 | PE100 | 196849539 | 27469032 | 4.55 | 6620 +/- 2722 |
| **Total Coverage** |  |  |  |  |  | **195.29X** |  |

| **Library** | **Protocol** | **Insert size (bp)** | **Read Length** | **Number of raw reads** | **Number of processed reads** | **Read Coverage** | **Estimated Insert size +/- sd (bp)** |
| --- | --- | --- | --- | --- | --- | --- | --- |
| Ppelagicus 300bp_V1 | Nextera | 160 | PE100 | 219872249 | 171327630 | 28.11 | 133 +/- 13 |
| Ppelagicus 500bp_V1 | Truseq nano | 380 | PE100 | 175198989 | 198082335 | 32.50 | 343 +/- 37 |
| Ppelagicus 600bp_V1 | Truseq nano | 480 | PE100 | 202270914 | 191709906 | 31.45 | 396 +/- 66 |
| Ppelagicus 850bp_V1 | Truseq nano | 730 | PE100 | 193571846 | 189459169 | 31.08 | 591 +/- 120 |
| Ppelagicus 2kb_V1 | Nextera Mate-pair | 2000 | PE100 | 233757470 | 25797170 | 4.23 | 1843 +/- 234 |
| Ppelagicus 3kb_V1 | Nextera Mate-pair | 2000 | PE100 | 191169907 | 58819331 | 9.65 | 2732 +/- 314 |
| Ppelagicus 4kb_V1 | Nextera Mate-pair | 2000 | PE100 | 201794028 | 67442086 | 11.07 | 3670 +/- 507 |
| Ppelagicus 6kb_V1 | Nextera Mate-pair | 2000 | PE100 | 210809550 | 46390405 | 7.61 | 5183 +/- 928 |
| **Total coverage** |  |  |  |  |  | **155.70X** |  |

**Supplementary Table 3. See Excel file**

**Supplementary Table 4.** Functional profiling of proteins with function-altering variants (PROVEAN score < −5)

a. Functional profiling of proteins with function-altering variants in *P. harrisi*.
| Term | ID | Name | Term size | Query size | Overlap size | P value |
| --- | --- | --- | --- | --- | --- | --- |
| 1 | HP:0000340 | Sloping forehead | 76 | 1101 | 18 | 0.0267 |
| 2 | HP:0006706 | Cystic liver disease | 18 | 1101 | 8 | 0.0331 |
| 3 | HP:0001197 | Abnormality of prenatal development or birth | 269 | 1101 | 48 | 0.000207 |
| 4 | HP:0001622 | Premature birth | 65 | 1101 | 16 | 0.0435 |
| 5 | HP:0001780 | Abnormality of toe | 285 | 1101 | 44 | 0.0298 |
| 6 | HP:0010297 | Bifid tongue | 14 | 1101 | 8 | 0.00307 |
| 7 | HP:0001830 | Postaxial foot polydactyly | 33 | 1101 | 11 | 0.0313 |
| 8 | HP:0000007 | Autosomal recessive inheritance | 1477 | 1101 | 171 | 0.00142 |
| 9 | HP:0001395 | Hepatic fibrosis | 57 | 1101 | 17 | 0.00154 |
| 10 | HP:0010936 | Abnormality of the lower urinary tract | 266 | 1101 | 43 | 0.0117 |
| 11 | HP:0000795 | Abnormality of the urethra | 181 | 1101 | 37 | 0.000154 |
| 12 | HP:0100627 | Displacement of the external urethral meatus | 156 | 1101 | 29 | 0.0211 |
| 13 | HP:0001159 | Syndactyly | 175 | 1101 | 33 | 0.00443 |
| 14 | HP:0006101 | Finger syndactyly | 120 | 1101 | 27 | 0.000964 |
| 15 | HP:0001177 | Preaxial hand polydactyly | 38 | 1101 | 12 | 0.026 |
| 16 | HP:0009142 | Duplication of bones involving the upper extremities | 119 | 1101 | 25 | 0.00858 |
| 17 | HP:0004275 | Duplication of hand bones | 119 | 1101 | 25 | 0.00858 |
| 18 | HP:0009997 | Duplication of phalanx of hand | 118 | 1101 | 24 | 0.0225 |
| 19 | HP:0000238 | Hydrocephalus | 169 | 1101 | 30 | 0.0383 |

b. Functional profiling of proteins with function-altering variants in flighted cormorants compared to *P. harrisi*
| Term | ID | Name | Term size | Query size | Overlap size | P value |
| --- | --- | --- | --- | --- | --- | --- |
| 1 | HP:0002242 | Abnormality of the intestine | 352 | 1058 | 54 | 0.0015 |
| 2 | HP:0012211 | Abnormal renal physiology | 284 | 1058 | 43 | 0.0225 |
| 3 | HP:0000083 | Renal insufficiency | 164 | 1058 | 34 | 0.00013 |

c. Functional profiling of function-altering variants in *P. pelagicus*
| Term | ID | Name | Term size | Query size | Overlap size | P value |
| --- | --- | --- | --- | --- | --- | --- |
| 1 | HP:0002202 | Pleural effusion | 11 | 875 | 6 | 0.0223 |
| 2 | HP:0001952 | Abnormal glucose tolerance | 16 | 875 | 7 | 0.0284 |
| 3 | HP:0040130 | Abnormal serum iron | 6 | 875 | 5 | 0.00558 |
| 4 | HP:0003452 | Increased serum iron | 6 | 875 | 5 | 0.00558 |
| 5 | HP:0001394 | Cirrhosis | 83 | 875 | 17 | 0.0185 |

**Supplementary Table 5.** List of the proteins with the strongest function-altering variants in *P. harrisi*.

| Ensembl chicken ortholog | Protein name | Variant | PROVEAN score | Preliminary Analysis | Sanger sequencing |
| --- | --- | --- | --- | --- | --- |
| ENSGALP00000002789 | **cut-like homeobox 1** | Q1120_S1123del | -15.704 | OK | Confirmed |
| ENSGALP00000005392 | family with sequence similarity 160, member A2 | A570_R588del<br>P602_A701del | -12.479<br>-154.202 | Annotation issue | N.D.<br>N.D. |
| ENSGALP00000006693 | torsin family 1, member B-like precursor | A222del | -12.610 | OK (§) frameshift | Confirmed |
| ENSGALP00000007432 | ankyrin repeat domain-containing protein 27 | L389del | -12.702 | OK | Confirmed |
| ENSGALP00000010467 | angiomin like 2 | L213_K219del<br>K236_L237insVSKWQQ | -14.537<br>-12.761 | OK<br>Q214_L220del (*)<br>-14.537 | Confirmed |
| ENSGALP00000011055 | Gallus gallus diaphanous homolog 1 (Drosophila) | E532_G533insVIK...IFQ<br>P571_P572insPPP...GAI | -14.083<br>-21.334 | Annotation issue | N.D. |
| ENSGALP00000011338 | RNA-binding protein 7 | P109_P110insARG...YK<br>Q | -18.483 | Annotation issue | N.D. |
| ENSGALP00000013107 | neurofilament, heavy polypeptide | K459_V534del<br>K672_E690del | -88.025<br>-17.752 | Annotation issue | N.D. |
| ENSGALP00000016953 | fasciculation and elongation protein zeta 1 | E279_R281del | -12.614 | OK | No PCR band.<br>GC content > 65% |
| ENSGALP00000018335 | TBC1 domain family, member 5 | M537_G544del | -34.202 | Annotation issue | N.D. |
| ENSGALP00000020006 | MICAL-like protein | E506_D508del | -13.397 | OK | Confirmed |
| ENSGALP00000020593 | ADNP homeobox 2 | I4_K36del | -123.241 | Assembly issue | Disproved |
| ENSGALP00000022179 | zinc finger protein 516 | G126_L130del | -14.716 | OK | No PCR band.<br>GC content >50% |
| ENSGALP00000022326 | KIAA1211 | E278_E284del | -19.786 | Annotation issue | N.D. |
| ENSGALP00000022477 | FAM210A | M228_K238del | -23.812 | OK | Disproved |
| ENSGALP00000023052 | NEDD4 binding protein 2 | S908_N910del | -15.178 | OK | Confirmed |
| ENSGALP00000026776 | zinc finger protein 395 | G44_A56del | -13.343 | Annotation issue | N.D. |
| ENSGALP00000036506 | **lectin, galactoside-binding, soluble, 3** | G45_P51del | -26.319 | OK | Confirmed |
| ENSGALP00000040904 | family with sequence similarity 160, member B2 | D142_L146del | -23.244 | OK | No PCR band.<br>GC content >65% |
| ENSGALP00000040933 | zinc finger protein 774 | K53del | -12.780 | OK | No PCR band.<br>GC content > 65% |
| ENSGALP00000042187 | R-spondin 2 | C21_R64del | -200.105 | Assembly issue | Disproved |
| ENSGALP00000042781 | BAH domain and coiled-coil containing 1 | V1152_E1167del | -16.307 | Annotation issue | N.D. |
| ENSGALP00000042911 | ankyrin repeat domain-containing protein 27 | L389del | -12.702 | OK | Confirmed |

Proteins carrying variants PROVEAN score < -12.5 in *P. harrisi*. Variants that were not caused by annotation issues were Sanger sequenced. (§) The mutation in Torsin family 1, member B-like precursor causes a frameshift and not a deletion. (\*) The variant in Angiomin like 2 is a 7aa deletion.

**Supplementary Table 6.** Functional profiling of CUX1 target genes in Hs578t cells.

|  | ID | Name | Term size | Query size | Overlap size | P value |
| --- | --- | --- | --- | --- | --- | --- |
| 1 | REAC:199977 | ER to Golgi Transport | 56 | 3356 | 22 | 0.042 |
| 2 | REAC:204005 | COPII (Coat Protein 2) Mediated Vesicle Transport | 56 | 3356 | 22 | 0.042 |
| 3 | REAC:0000000 | Reactome pathways | 6634 | 3356 | 1341 | 1.39E-07 |
| 4 | REAC:392499 | Metabolism of proteins | 731 | 3356 | 170 | 0.0328 |
| 5 | REAC:74160 | Gene Expression | 1078 | 3356 | 261 | 1.35E-05 |
| 6 | REAC:1430728 | Metabolism | 1351 | 3356 | 296 | 0.0156 |
| 7 | REAC:1640170 | Cell Cycle | 456 | 3356 | 126 | 3.99E-05 |
| 8 | REAC:69278 | Cell Cycle | 378 | 3356 | 106 | 0.000182 |
| 9 | REAC:453279 | Mitotic G1-G1/S phases | 106 | 3356 | 36 | 0.016 |
| 10 | REAC:1852241 | Organelle biogenesis and maintenance | 268 | 3356 | 80 | 0.00031 |
| 11 | REAC:5617833 | **Assembly of the primary cilium** | 152 | 3356 | 53 | 0.00012 |
| 12 | REAC:5620924 | **Intraflagellar transport** | 37 | 3356 | 18 | 0.0057 |
| 13 | REAC:2559583 | Cellular Senescence | 155 | 3356 | 47 | 0.0327 |

**Supplementary Table 7.** Test for positive selection in previously identified cilia related genes carrying function-altering mutations in *P. harrisi*.

| Gene | *P. harrisi*<br>branch<br>($\omega$) | Background<br>Phylogeny<br>($\omega$) | Log-<br>likelihood<br>(branch<br>model) | Log-<br>likelihood<br>($M_0$) | P-value ($\chi^2$ test) |
| --- | --- | --- | --- | --- | --- |
| *Ofd1* | 1.92 | 0.33 | -34752.80522 | -34754.57355 | 0.06005 |
| *Talpid3* | 0.35 | 0.37 | -61668.50971 | -61668.51784 | 0.898719 |
| *Evc* | 1.93 | 0.34 | -31125.50247 | -31127.18311 | 0.065196 |
| *Wdr34* | 0.37 | 0.24 | -20893.5747 | -20893.67402 | 0.655853 |
| ***Gli2*** | **1.10** | **0.11** | **-39721.49349** | **-39726.09523** | **0.002416 (\*\*)** |
| *Gli3* (†) | 0.04 | 0.15 | -33534.29128 | -33535.53804 | 0.114354 |
| *Fat1* | 0.17 | 0.11 | -115981.2078 | -115981.8595 | 0.253611 |
| *Dchs1* | 0.18 | 0.09 | -93197.72901 | -93198.54569 | 0.201247 |
| *Dvl1* | 0.20 | 0.07 | -16807.654446 | -16807.934377 | 0.454341 |
| *Ift122* | 0.72 | 0.07 | -20435.710607 | -20437.573224 | 0.053604 |
| *Dync2h1* | 0.17 | 0.11 | -54031.742919 | -54032.027581 | 0.450537 |
| *Kif7* | 0.09 | 0.12 | -43536.031228 | -43536.155293 | 0.618416 |

The ratio of non-synonymous to synonymous substitutions ( $\omega = dN/dS$ ) was estimated using a branch likelihood model. $\omega < 1$ indicates purifying selection, $\omega > 1$ indicates positive selection. (†) *Gli3* serves as a control for its paralog *Gli2*.

**Supplementary Table 8.**
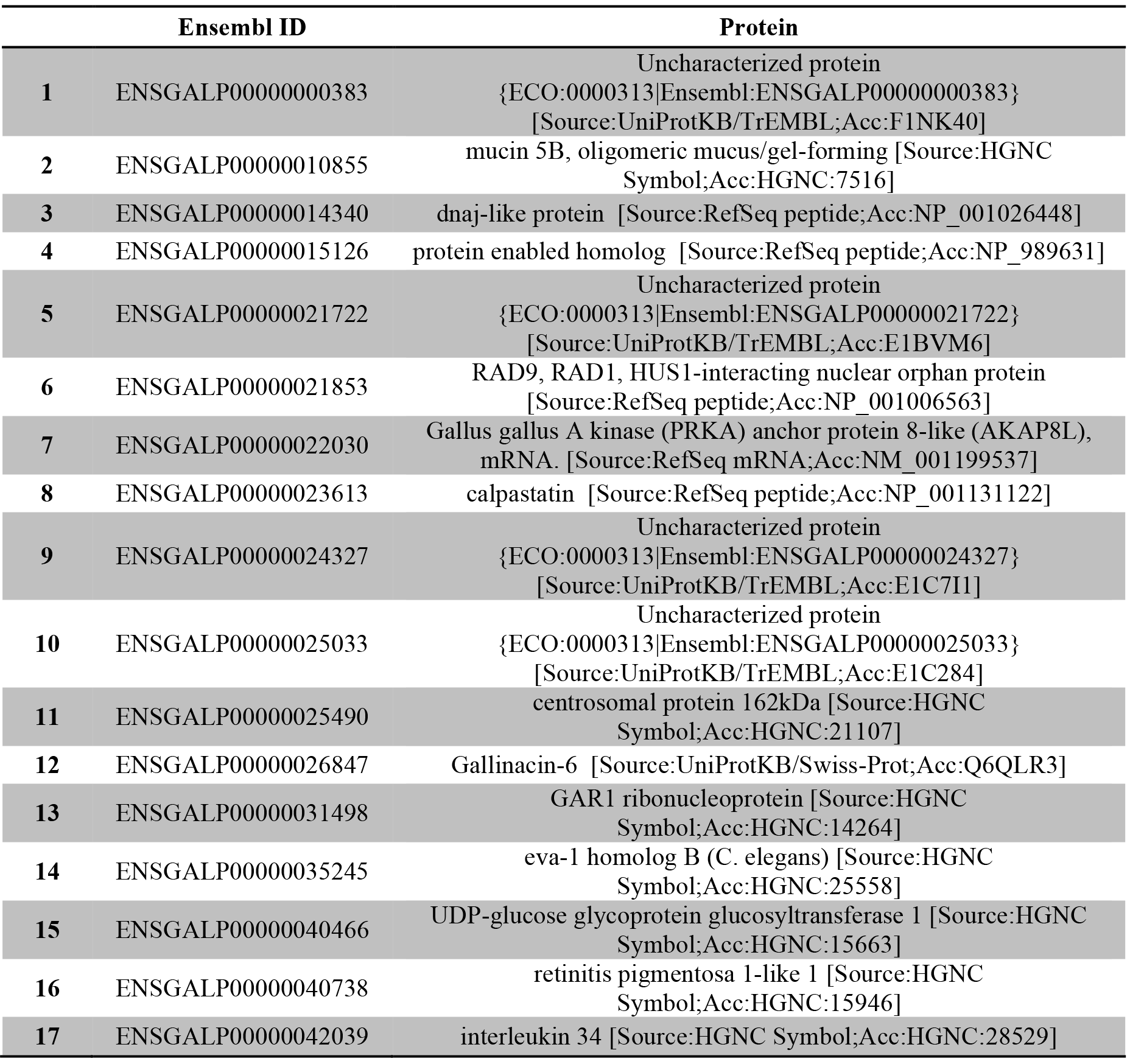
List of genes not annotated in *P. harrisi* but annotated in *P. auritus, P. brasilianus* and *P.pelagicus*.

**Supplementary Table 9.**
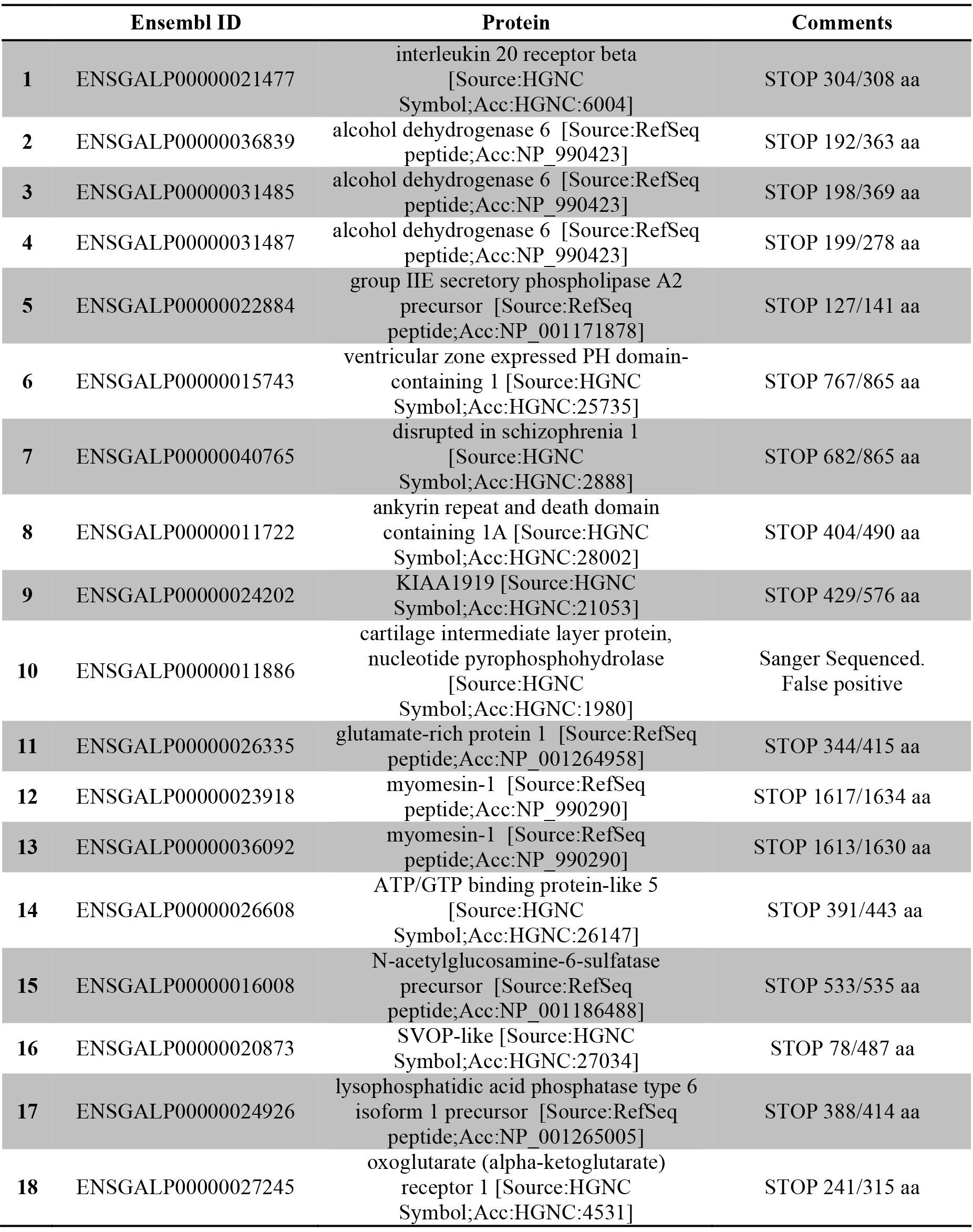
Curated list of proteins with putative premature stop codons in *P. harrisi* that are absent in in *P. auritus, P. brasilianus* and *P.pelagicus*.

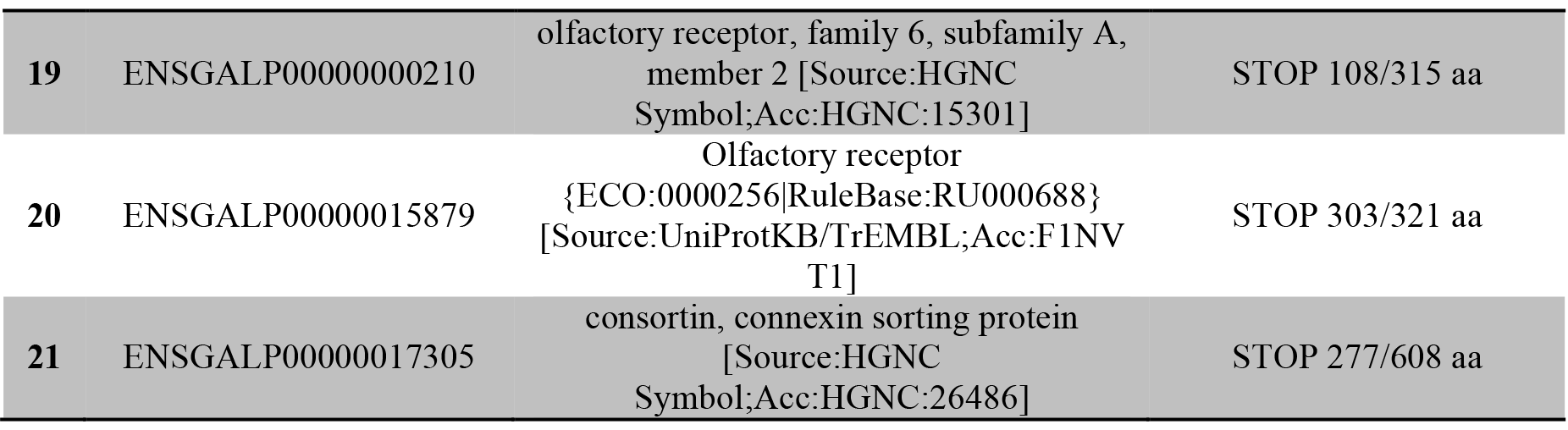

**Supplementary Table 10.**
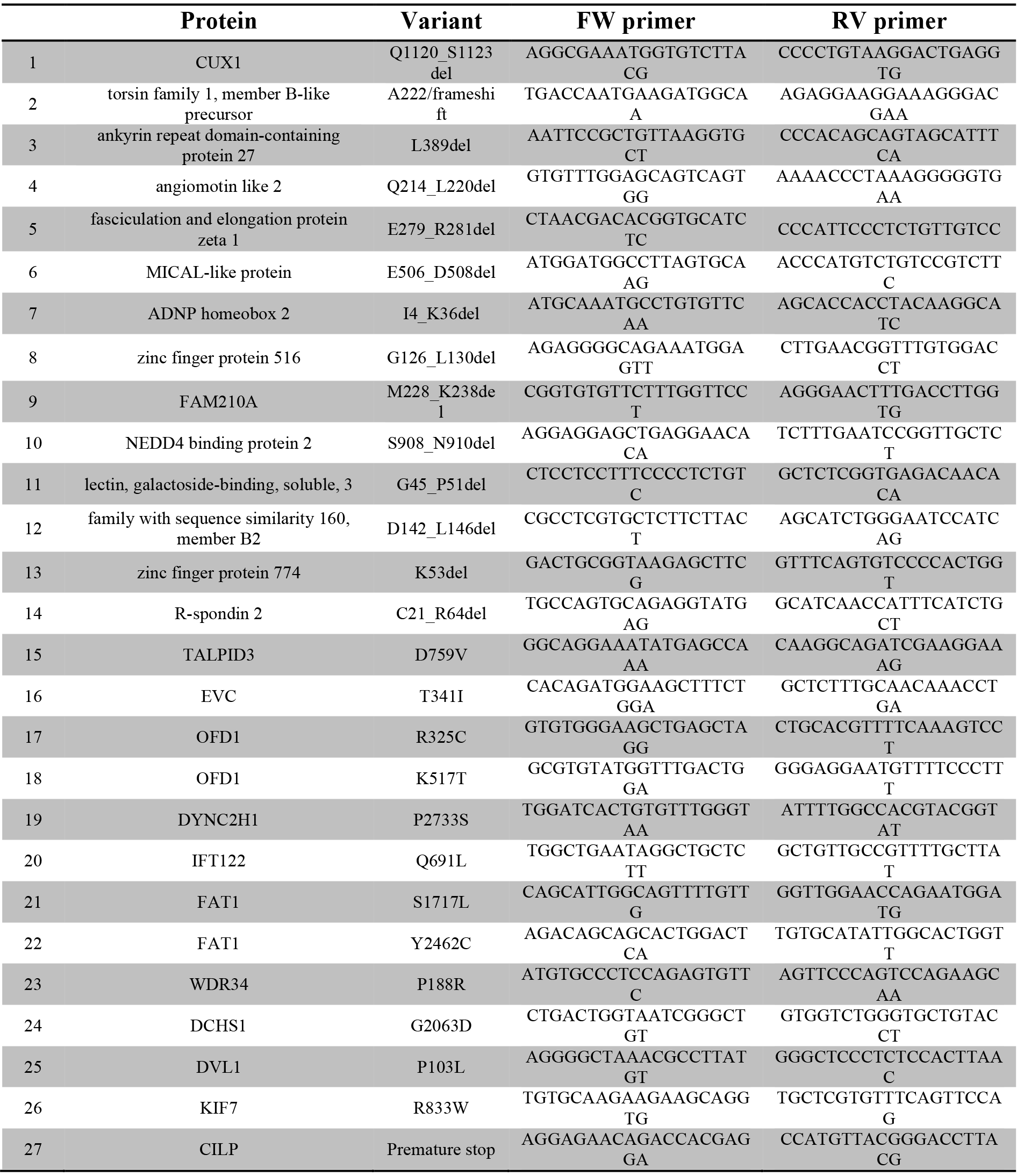
Primers used to verify predicted variants in the sequenced cormorants.

**Supplementary Table 11.**
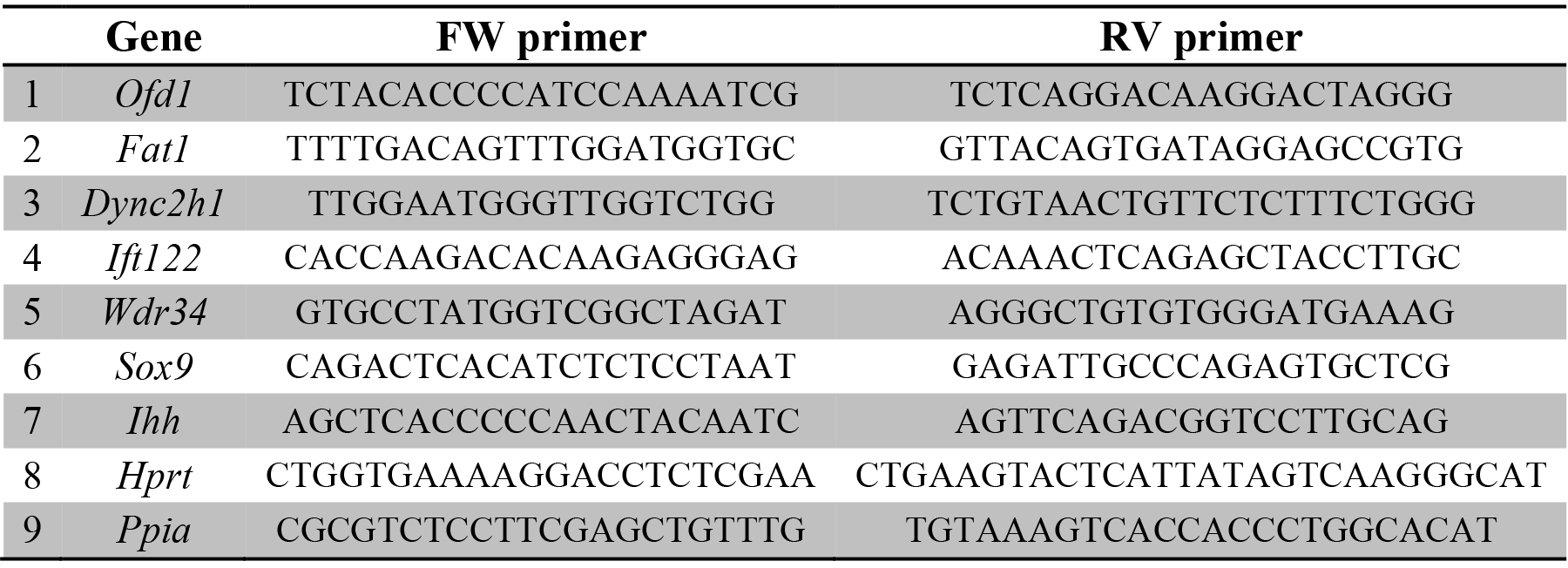
Primers used for quantitative real time PCR in ATDC5 cell lines.

